# Single-cell heterogeneity in the chloroplast redox state mediates acclimation to stress in a marine diatom

**DOI:** 10.1101/319517

**Authors:** Avia Mizrachi, Shiri Graff van Creveld, Orr H. Shapiro, Shilo Rosenwasser, Assaf Vardi

## Abstract

Diatoms are photosynthetic microorganisms of great ecological and biogeochemical importance, forming vast blooms in diverse aquatic ecosystems. Current understanding of phytoplankton acclimation to stress is based on population-level analysis, masking cell-to-cell variability. Here we investigated heterogeneity within *Phaeodactylum tricornutum* populations in response to oxidative stress, which is induced by environmental stress conditions. We combined flow cytometry and a microfluidics system for live imaging to measure redox dynamics at the single-cell level using the roGFP sensor. Chloroplast-targeted roGFP exhibited a light-dependent, bi-stable oxidation pattern in response to H_2_O_2_, revealing distinct subpopulations of sensitive oxidized cells and resilient reduced cells. Subpopulation proportions depended on growth phase, linking the bi-stable phenotype to proliferation. Oxidation of chloroplast-targeted roGFP preceded commitment to cell death and was used as a novel cell fate predictor. We propose that light-dependent metabolic heterogeneity results in differential stress responses that regulate cell fate within diatom populations.

## Introduction

Diatoms are considered amongst the most successful and diverse eukaryotic phytoplankton groups, and are estimated to contribute 20% of global net primary production^1–3^. They form massive blooms and are thus central to the biogeochemical cycling of important elements such as carbon, nitrogen, phosphate, iron and silica, in addition to their important role at the base of marine food webs^1,2,4–6^. As other phytoplankton, diatoms need to constantly acclimate to physicochemical gradients in a fluctuating environment. They are exposed to stress from different biotic and abiotic origins such as grazing, viruses, bacteria, allelopathic interactions, light availability, and nutrient limitations^7–14^. Importantly, induction of programmed cell death (PCD) in response to different stressors has been suggested as an important mechanism contributing to the fast turnover of phytoplankton and the rapid bloom demise^7,8,15^.

Recent studies suggested that diatoms can differentially response to diverse environmental cues based on compartmentalized redox fluctuations that also mediate stress-induced PCD^16–18^. Reactive oxygen species (ROS) are known to play an important role in sensing stress and additional signals across kingdoms, from bacteria to plants and animals^19–23^. They are produced as byproducts of oxygen-based metabolism in respiration and photosynthesis, by ROS generating enzymes, and due to various stress conditions^16,22,24–27^. To maintain redox balance and avoid oxidative damage, cells harbor various ROS scavenging enzymes and small antioxidant molecules that regulate and buffer ROS levels, such as glutathione (GSH), ascorbate and NADPH. ROS can cause fast post-translational modifications of proteins through oxidation, affecting their activity prior to changes in gene expression^22^. The specificity of the ROS signal is derived from the specific chemical species, its concentration, sub-cellular localization, temporal dynamics, and available downstream ROS-sensitive targets^16,22,24,28–30^. Therefore, ROS production and redox metabolic networks can be used to sense and integrate information of both the metabolic state of the cell and its microenvironment.

H_2_O_2_ is a relatively mild and stable ROS that can accumulate in cells due to various stress conditions, thus often serves as a signaling molecule^19–23, 31^. It has a preferential activity towards cysteine residues, and can remodel the redox-sensitive proteome network^17,23,28^. In addition, H_2_O_2_ can diffuse across membranes (depending on membrane properties) and through aquaporins channels^32^. Combined properties as lower toxicity, diffusibility and selective reactivity make H_2_O_2_ suitable for studying signaling in various biological systems^23,28^. Since many environmental stressors induce ROS generation, application of H_2_O_2_ can reproduce the downstream cellular response. H_2_O_2_ application in marine diatoms led to oxidation patterns similar to other environmental stressors^16,17^. It also led to the induction of cell death, in a dose-dependent manner, with characteristics of PCD that included externalization of phosphatidylserine, DNA laddering, and compromised cell membrane^16^. In addition, early oxidation of the mitochondrial GSH pool preceded subsequent cell death at the population level following exposure to H_2_O_2_ and diatom-derived infochemicals in the model diatom *Phaeodactylum tricornutum*^16,33^.

In the current work, we investigated phenotypic variability within diatom populations in response to oxidative stress. Until recently, research of phytoplankton’s responses to environmental stress was carried out primarily at the population level, masking heterogeneity at the single-cell level. Cell-to-cell variability could result in different cellular strategies employed by the population. We established single-cell approaches to measure *in vivo* oxidation dynamics in the model diatom *P. tricornutum* using flow cytometry and microfluidics live-imaging. To measure oxidation dynamics of specific organelles, we used *P. tricornutum* strains expressing redox-sensitive GFP (roGFP) targeted to different sub-cellular compartments^17^. The oxidation of roGFP is reversible, and can be quantified using ratiometric fluorescence measurements^34^. The roGFP oxidation degree (OxD) reports the redox potential of the GSH pool (E_GSH_), which represents the balance between GSH and its oxidized form^34^. Therefore, roGFP OxD provides a metabolic input of the redox state of the cell, and represents the oxidation state of native proteins in the monitored organelle^17,34^. By measuring redox dynamics at single-cell resolution, we uncovered a previously uncharacterized phenotypic heterogeneity in the response of a marine diatom to oxidative stress. We revealed a bi-stable response in the chloroplast E_GSH_ following H_2_O_2_ treatment, exposing distinct subpopulations with different cell fates. Our results revealed a specific link between oxidation patterns in the chloroplast and cell fate regulation.

## Results

### Bi-stable chloroplast roGFP oxidation in response to oxidative stress reveals distinct subpopulations

To assess roGFP oxidation in an organelle-specific manner, we measured OxD in *P. tricornutum* strains expressing roGFP targeted to the chloroplast, nucleus and mitochondria using flow cytometry. At steady-state conditions without perturbations, the OxD distribution in the population had a single distinct peak, representing a reduced state at all examined compartments (Fig. 1 A, E, I and Figs. S1, S2). Application of H_2_O_2_ led to organelle-specific and dose-dependent oxidation patterns in these organelles (Fig. 1).

**Fig. 1.**
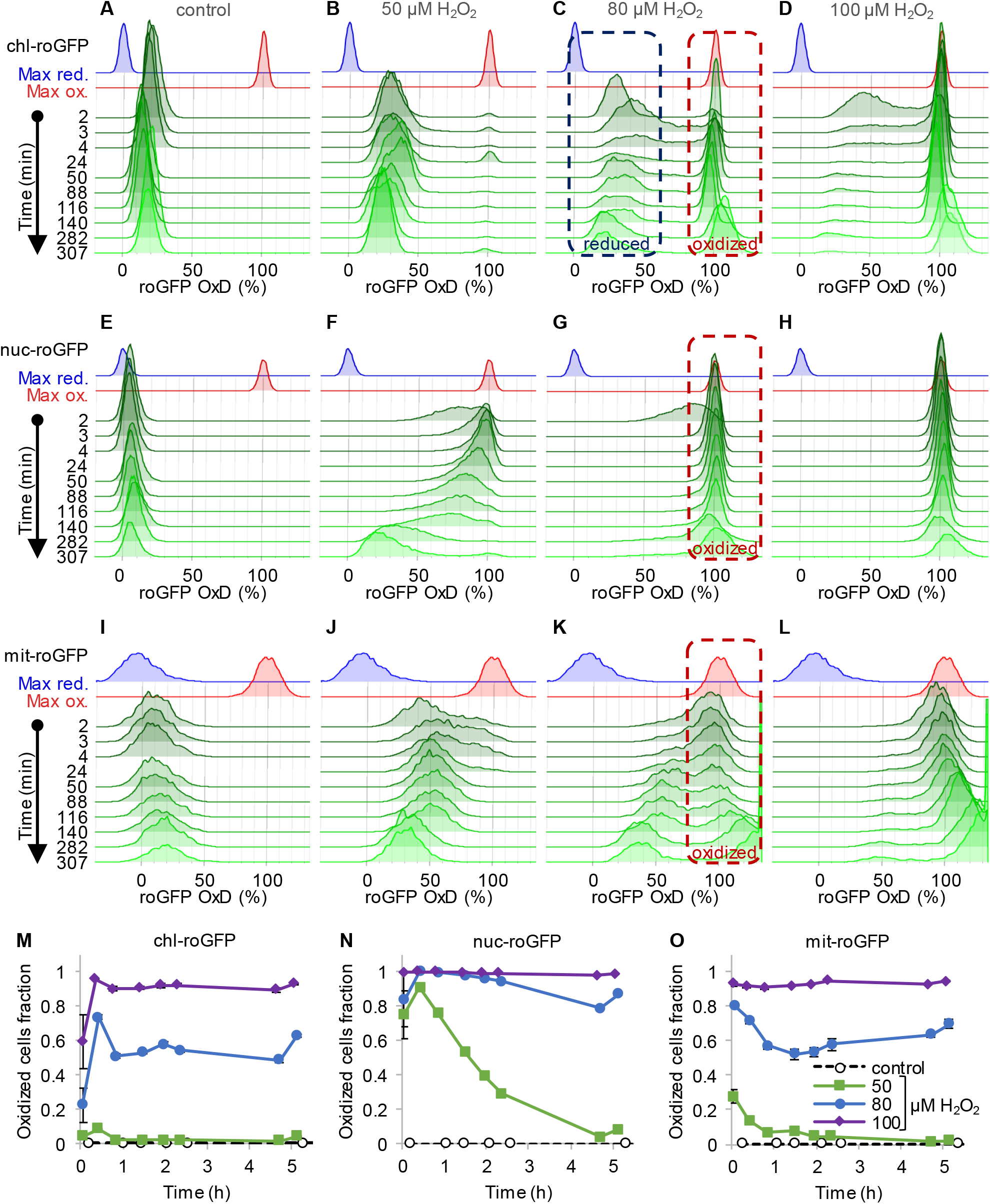
Oxidation of roGFP in response to H_2_O_2_ is organelle-specific and reveals heterogeneity at the single-cell level. The distribution of roGFP OxD in the population over time was measured by flow cytometry in *P. tricornutum* cells expressing roGFP targeted to the chloroplast (chl-roGFP, A-D, M), nucleus (nuc-roGFP, E-H, N) and mitochondria (mit-roGFP, I-L, O). (***A-L***) Oxidation of roGFP in response to 0 μM (A,E,I), 50 μM (B,F,J), 80 μM (C,G,K), and 100 μM (D,H,L) H_2_O_2_. Maximum reduction (blue) and oxidation (red) of roGFP following additions of 2 mM DTT or 200 μM H_2_O_2_ respectively are shown as a reference. The “oxidized” and “reduced” subpopulations are marked by red and blue dashed boxes respectively (C, G, K). The experiment was done in triplicates, for visualization the first replica is shown except for the first 4 min in which all replicates are shown for higher temporal resolution. Each histogram consists of >8000 (A-D), >5900 (E-H) and >1400 (I-L) roGFP-positive cells. (***M-O***) The fraction of the “oxidized” subpopulation over time upon exposure to 0-100 μM H_2_O_2_. Mean ± SEM, n=3 biological repeats. SEM lower than 0.018 are not shown.

The chloroplast-targeted roGFP (chl-roGFP) exhibited a distinct bi-stable response following treatments of 50-100 μM H_2_O_2_, revealing two distinct subpopulations of “oxidized” and “reduced” cells (Fig. 1 B-D and Fig. S3 A-C). These subpopulations emerged within the first few minutes after H_2_O_2_ addition (Fig. 1 B-D). The existence of these subpopulation is masked in bulk analysis, demonstrating the importance of single-cell measurements (Fig. S1). In the “oxidized” subpopulation, roGFP completely oxidized (∼100%) in response to H_2_O_2_, reaching a similar distribution of the fully oxidized positive control (200 μM H_2_O_2_) (Fig. 1 B-D and Fig. S3 G). However, in the “reduced” subpopulation roGFP reached lower values of 30-43% OxD within 2-24 minutes post treatments, and then gradually recovered (Fig. 1 B-D and Fig. S3 G). Only a minor fraction of the cells displayed intermediate oxidation, suggesting that these subpopulations represent discrete redox states. Interestingly, a larger fraction of cells was within the “oxidized” subpopulation at 20-25 minutes post treatment compared to later time points, indicating that some cells were able to recover during this time (Fig. 1 M). The proportion between these subpopulations stabilized after 46-51 min post treatment, and was H_2_O_2_-dose dependent, as more cells were within the “oxidized” subpopulation at higher H_2_O_2_ concentrations (Fig. 1 A-D, M). The quick emergence of stable co-existing “oxidized” and “reduced” subpopulations exposed underlying heterogeneity within the diatom population, resulting in a differential response to oxidative stress.

This clear bi-stable pattern was unique to the chloroplast-targeted roGFP. Nuclear roGFP displayed a continuous distribution in response to H_2_O_2_ treatments, and no distinct subpopulations could be observed (Fig. 1 F-H). Within minutes post treatment, nuclear roGFP exhibited fast oxidation even in response to a low H_2_O_2_ concentration of 50 μM, which had only a mild effect on the chloroplast (Fig. 1 B, F). At that concentration, nuclear roGFP oxidation was followed by a gradual and much slower recovery, which lasted >5 hours post treatment (Fig. 1 F). At higher concentrations, the entire population was oxidized within 3 minutes post treatment, and most cells remained stably oxidized >5 hours post treatment (Fig. 1 G-H). The mitochondrial roGFP exhibited a heterogeneous redox response, as seen in the 80 μM and 100 μM H_2_O_2_ treatments starting at ∼24 minutes post treatment (Fig. 1 K-L). However, distinct subpopulations were not clearly separated until later stages, and were not detected consistently in different experiments (Fig. 1 K-L and Fig. S4). Therefore, we chose to focus on the chl-roGFP strain, which revealed two discrete subpopulations.

### Oxidation of chl-roGFP precedes the induction of cell death

Next, we examined the possible link between early chloroplast E_GSH_ oxidation and subsequent mortality. We quantified cell death 24 hours post H_2_O_2_ treatment using flow cytometry measurements of Sytox green staining, which selectively stains nuclei of dying cells. The fraction of “oxidized” cells 1-2 h post treatment was correlated with the fraction of dead cells at 24 h (Fig. 2 A, Spearman’s correlation coefficient r_S_=0.98, *P*=4.6·10^−8^), suggesting that early oxidation in the chloroplast in distinct subpopulations may predict cell fate at much later stages.

**Fig. 2.**
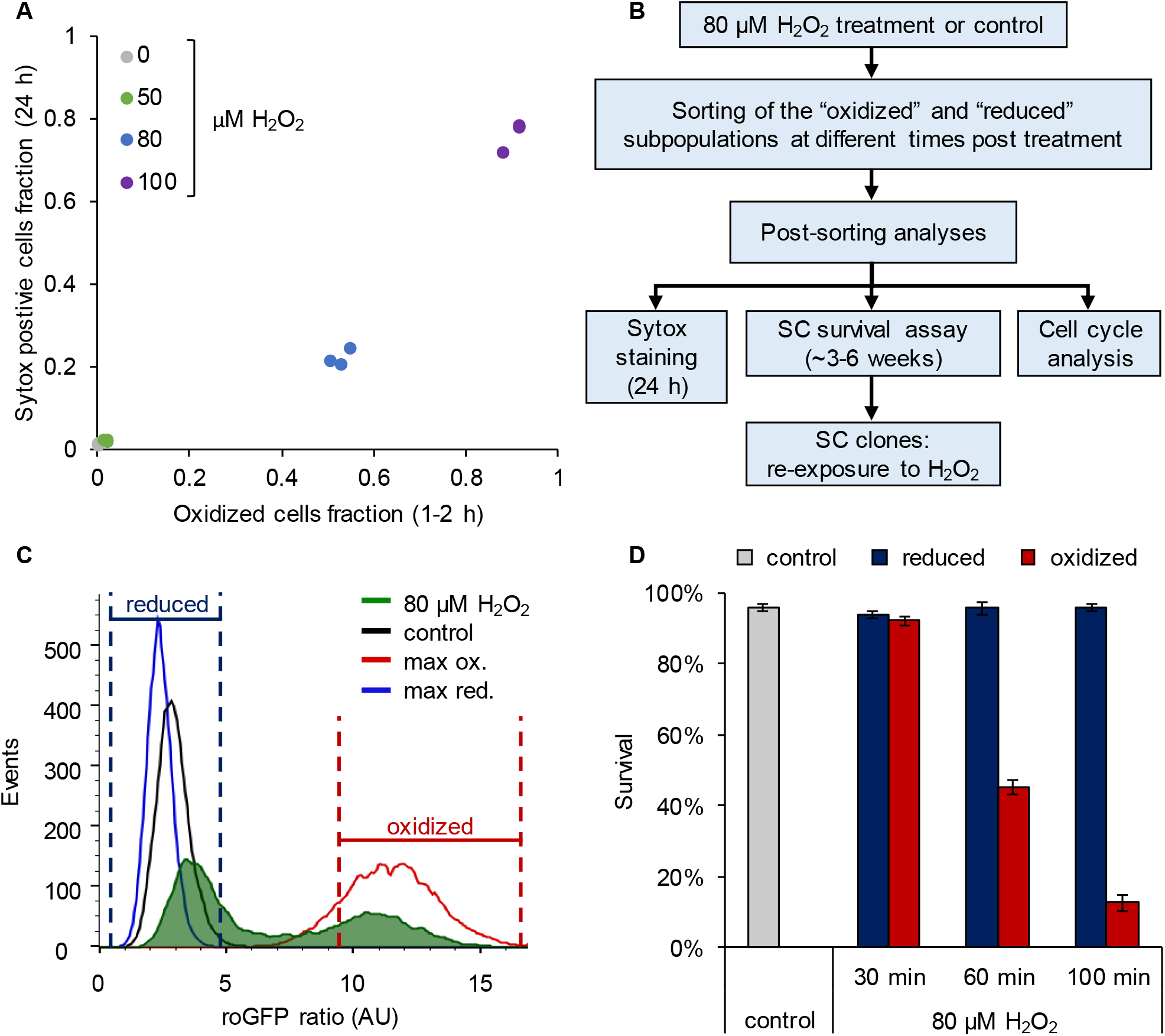
Early oxidation of chl-roGFP in a subpopulation precedes the induction of cell death. (***A***) The fraction of dead cells measured by Sytox positive staining 24 h post 0-100 H_2_O_2_ treatment as a function of the fraction of chl-roGFP “oxidized” cells 1-2 h post treatment (see Fig. 1C). N=3 biological repeats per treatment, individual samples are shown. (***B***) Schematic representation of sorting experiments layout and post-sorting analyses. SC – single-cell. (***C***) Representative histograms of chl-roGFP oxidation distribution measured by roGFP ratio (arbitrary units, AU, see methods) 100 min following 80 μM H_2_O_2_ treatment (green). Maximum oxidation (200 μM H_2_O_2_, red), maximum reduction (2 mM DTT, blue), and untreated control (black) are shown for reference. Gates used for sorting the “oxidized” and “reduced” subpopulations based on chl-roGFP oxidation are shown, untreated control cells were sorted based on positive roGFP fluorescence. (***D***) Survival of individual cells that were sorted into fresh media and regrown, measured by the number of colony forming units divided by the number of sorted cells. Data is shown as mean ± SEM, n≥6 biological repeats, each of 24-48 single cells per time-point per subpopulation that were sorted into separate wells.

To investigate directly the link between early chl-roGFP oxidation and subsequent cell death, we used fluorescence-activated cell sorting (FACS) to sort cells based on chl-roGFP oxidation and measured their survival. Single cells of the “oxidized” and “reduced” subpopulations were sorted into fresh media at different time-points following the addition of 80 μM H_2_O_2_, and colony forming units were counted to assess survival (Fig. 2 B-D). When sorted 30 minutes post treatment, the “oxidized” subpopulation exhibited a high survival rate (92.3 ± 1.4%) that was similar to the “reduced” subpopulation (94.1 ± 1.1%, *P*=0.24, paired t-test) and to sorted untreated control (96 ± 0.9%, *P*=0.29, Dunnett test, Fig. 2 D). However, at later time-points the survival of the “oxidized” subpopulation gradually diminished, and was significantly lower than both the “reduced” and control sorted cells (*P*<0.001 for all comparisons, paired t-test for comparisons with “reduced” cells, Dunnett test for comparisons with control). When sorted 60 min post treatment almost half of the “oxidized” cells recovered (45.1 ± 2%), but when sorted 100 min following treatment only 12.7 ± 2.1% survived (*P*<0.001, Tukey test, Fig. 2 D). These results suggest that after a distinct exposure time, cell death is induced in an irreversible manner in the “oxidized” subpopulation. In contrast, the “reduced” subpopulation from the same culture and treatment exhibited a high survival rate similar to the control at all time-points examined, demonstrating its resilience to the stress (Fig. 2 D, *P*≥0.86 for all comparisons, Dunnett test). In agreement with these findings, cell death measurements using Sytox staining of sorted subpopulations also showed higher mortality in the “oxidized” cells compared to the “reduced” and the control cells, which remained viable (Fig. S5). Taken together, these results demonstrate that the “oxidized” subpopulation was sensitive to the oxidative stress applied, which led to induction of cell death in those cells, while the “reduced” subpopulation was able to survive. In addition, we detected a distinct phase of “pre-commitment” to cell death, ranging approximately 30-100 min in most cells, during which the fate of the “oxidized” subpopulation is still reversible upon removal of the stress.

### Early oxidation of chloroplast E_GSH_ predicts cell fate at the single-cell level

In order to track oxidation dynamics and subsequent cell fate of individual cells, we established a microfluidics platform for *in-*vivo long-term epifluorescence imaging, under controlled flow, light and temperature conditions adapted for diatom cells (Fig. 3, Fig. S6 and movies S1, S2). We introduced cells expressing chl-roGFP into a custom-made microfluidics device, let the cells settle, and introduced treatments of either 80 μM H_2_O_2_ or fresh media (control) continuously for 2.5-3 hours, after which the treatment was washed by fresh media (see methods). In addition, the use of microfluidics enabled imaging of the basal OxD state of single cells prior to treatment, as well as the introduction of Sytox green at the end of the experiment to visualize cell death. We detected the distinct “oxidized” and “reduced” subpopulations following 80 μM H_2_O_2_ treatment, similar to the flow cytometry experiments (Fig. 3 C, F, and movie S1). However, no clear differences were observed in their OxD prior to treatment (Fig. S7 and movie S1). The separation between the subpopulations emerged within 20 minutes of exposure to 80 μM H_2_O_2_, and remained stable over the course of the experiment with the “oxidized” subpopulation maintaining a high OxD above 80% (Fig. 3 F, Fig. S7 B and movie S1). The “reduced” subpopulation exhibited an immediate response to H_2_O_2_ comparable with flow cytometry measurements, from 25-45% OxD before treatment to 30-65% OxD during the first 20 minutes post 80 μM H_2_O_2_ treatment (Fig. 3 F, Fig. S7 B, and movie S1). Following this initial oxidation, the “reduced” cells recovered gradually over the next hours, reducing to 5-25% oxidation 8 hours post treatment, below the initial basal state (Fig. 3 F, movie S1). A gradual slow reduction was also observed in the control cells over the course of the experiment (Fig. 3 E, movie S2), which may represent acclimation to the experimental setup or a diurnal redox alteration. Control cells did not oxidize in response to addition of fresh media (Fig. 3 E and Fig. S7 A), excluding the possibility that the oxidation observed in 80 μM H_2_O_2_ treated cells was due to shear stress during treatment.

**Fig. 3.**
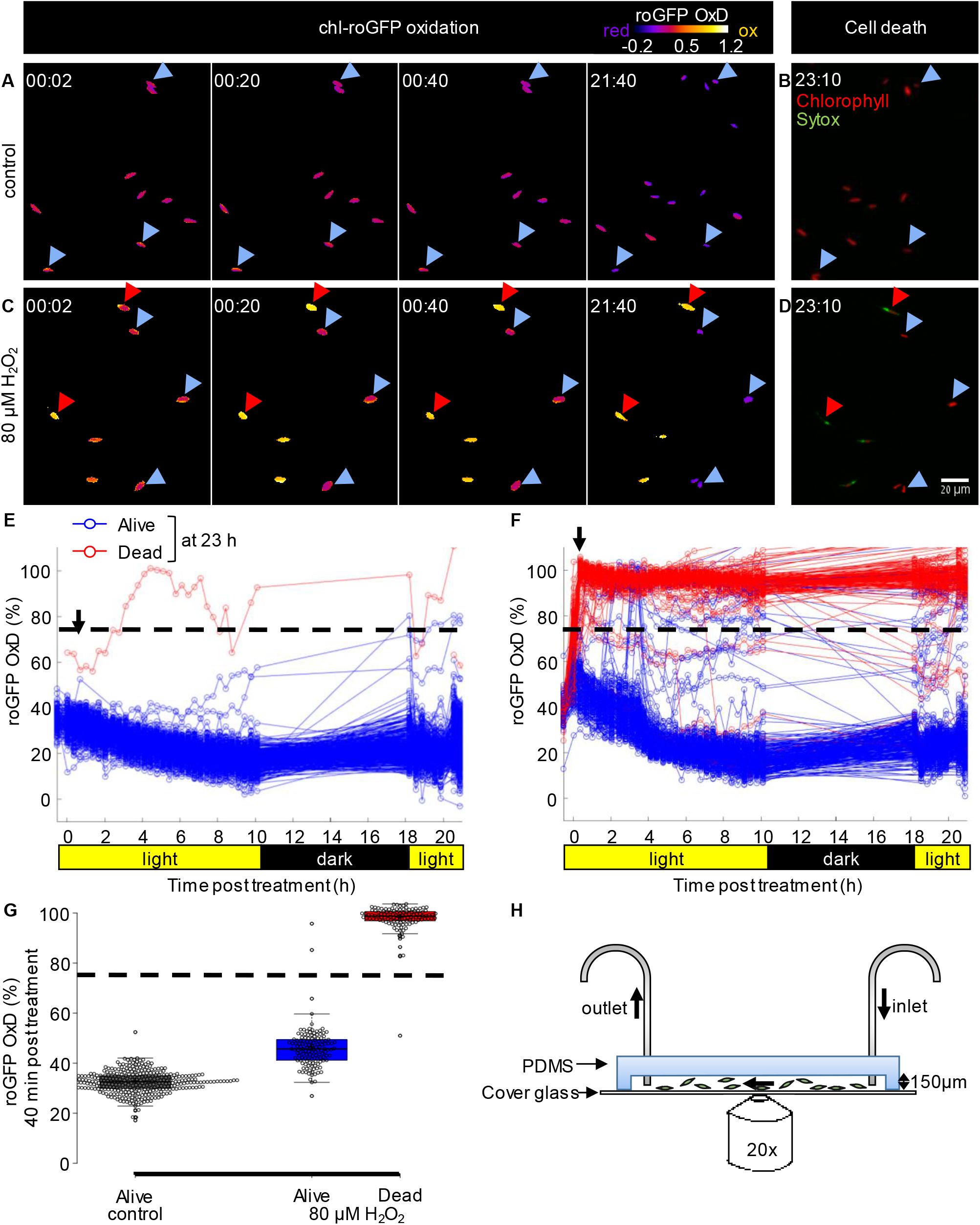
Tracking redox dynamics of individual cells using long term *in vivo* imaging in a microfluidics setup. Oxidation of chl-roGFP was imaged over time using a customized microfluidic setup and epifluorescence microscopy. Cells were imaged following treatment with either fresh media (control; A, B, E) or 80 μM H_2_O_2_ (C, D, F). To quantify cell death, cells were stained with Sytox ∼23 h post treatment. (***A, C***) Representative frames depicted in pseudo-color of calculated roGFP OxD at different times post treatment (hh:mm). Two subpopulations of “oxidized” (red arrows) and “reduced” (blue arrows) cells were detected in treated cells. (***B, D***) Overlay of Sytox staining (green, dead cells) and chlorophyll auto-fluorescence (red) at 23:10 h post treatment. (***E, F***) Quantification of chl-roGFP OxD per cell over time post treatment. Color is based on cell fate as measured ∼23 h post treatment: blue – alive (Sytox negative), red – dead (Sytox positive). OxD values above 110% likely result from auto-fluorescence leakage and are not shown. (***G***) Box-plot of chl-roGFP OxD 40 min post treatment (arrow in E, F) of cells that were grouped based on their cell fate as measured ∼23 h post treatment. Grey – control alive; blue – 80 μM H_2_O_2_ alive; red – 80 μM H_2_O_2_ dead. Circles - single cells; cross – mean; colored box – 1^st^ to 3^rd^ quartiles; horizontal line within the box – median. (***E-G***) N≥250 cells per treatment from ≥3 different fields of view. Horizontal dashed line represents the threshold used for cell fate predictions. (***H***) Side view of the microfluidics chip design.

We detected a clear correlation between initial oxidation in the chloroplast in response to oxidative stress and subsequent cell fate (Fig. 3 A-G). Cells that exhibited high chl-roGFP oxidation within the first 40 minutes also died at a much later stage, while cells that maintained a lower OxD were able to recover (Fig. 3 G). Logistic regression modelling of cell death as a function of chl-roGFP OxD at this time-point revealed a threshold of ∼74% OxD, that separated between cells that were dead or alive at the end of the experiment (Fig S8). Using this threshold, the OxD of chl-roGFP 40 minutes post H_2_O_2_ treatment predicted subsequent death of individual cells as measured ∼23 hours post treatment with a high accuracy of 98.8% (0.8% false positive, 1.7% false negative; Fig. 3 G and Fig. S8). These results corroborate the flow-cytometry analysis and demonstrate that under these conditions early chloroplast E_GSH_ response accurately predicts cell fate at the single-cell level.

### The distinct subpopulations derive from phenotypic variability and not from variable genetic backgrounds

The differential chloroplast oxidation of the observed subpopulations could be due to genetic variability or due to phenotypic plasticity within the population. To differentiate between the two scenarios, we sorted chl-roGFP individual cells of the “oxidized” and “reduced” subpopulations 30 and 100 minutes post 80 μM H_2_O_2_ treatment as well as untreated control cells, and regrew them to generate clonal populations derived from cells exhibiting specific phenotypes. The clonal progeny cultures were subsequently exposed to 80 μM H_2_O_2_ and their chl-roGFP oxidation was measured. The two distinct subpopulations were detected in all the clones measured, and the fraction of the “oxidized” subpopulation was again correlated with cell death (Fig. 4 and Fig. S9). Therefore, the different subpopulations observed did not originate from genetic differences, but rather represent phenotypic variability within isogenic populations.

**Fig. 4.**
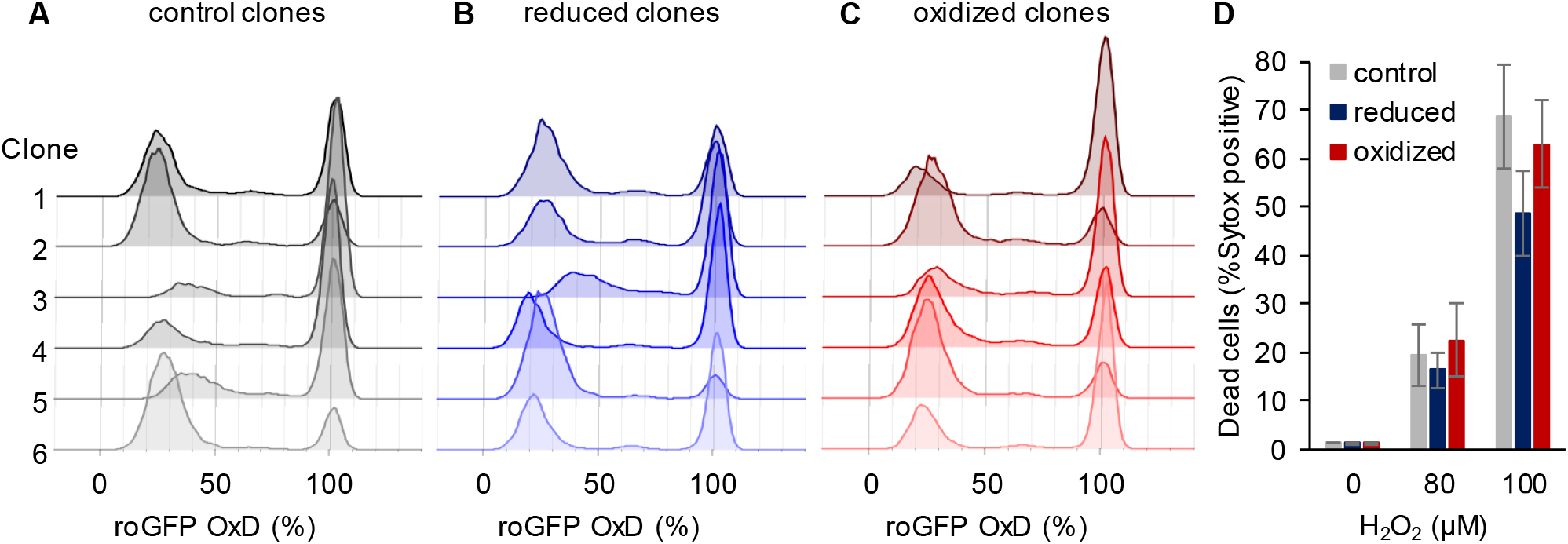
Clonal populations derived from sorted cells maintain the bi-stable phenotype of chl-roGFP response to H_2_O_2_. (***A-C***) The distribution of chl-roGFP OxD (%) 40-45 min post 80 μM H_2_O_2_ treatment in clonal populations derived from sorted single cells of different origins, 3 weeks post sorting. The “reduced” (B) and “oxidized” (C) subpopulations were sorted 30 minutes post 80 μM H_2_O_2_ treatment based on chl-roGFP oxidation (Fig. 2 C); “control” (A) – clones of untreated cells that were sorted based on positive roGFP fluorescence. Each histogram is of a single clone, ≥9900 cells per histogram, 6 representative clones per group are shown. (***D***) The fraction of dead cells 24 h post H_2_O_2_ treatment of the different clones shown in (A-C) as measured by positive Sytox staining. Data is shown as mean ± SEM, n=6 clones per group per treatment.

### The “oxidized” subpopulation is enriched with cells at G_1_ phase

One possible source for phenotypic variability in genetically homogenous populations can be explained by differences in the cell cycle phase, as the cell cycle is linked to metabolic changes including redox oscilations^35–38^. Therefore, we sorted the “oxidized” and “reduced” subpopulations 30 min following 80 μM H_2_O_2_ treatment into a fixation solution, and stained the fixed cells with 4′,6-diamidino-2-phenylindole (DAPI) to quantify DNA content for cell cycle analysis. Considering *P. tricornutum* divides about once a day^39^, we assumed that at this early time point the DNA content is likely to reflect the status of the cell prior to treatment, and is not yet affected by it. The sorted “oxidized” subpopulation had a higher fraction of cells at G_1_ (86.9 ± 1.8%) compared to control untreated cells (76.8 ± 0.7%, *P*=0.0024, Tukey test). The “reduced” subpopulation on the other hand had a smaller fraction of G_1_ cells (68.7 ± 2.2%) compared to both control (*P*=0.011, Tukey test) and “oxidized” cells (*P*=0.0001, paired t-test), and exhibited a larger fraction of G_2_/M cells (Fig. 5). These results demonstrate that although cell cycle phase alone cannot explain the differences between the subpopulations, it is linked to the chloroplast E_GSH_ response to oxidative stress and may represent an important factor that affects H_2_O_2_ sensitivity in the population.

**Fig. 5.**
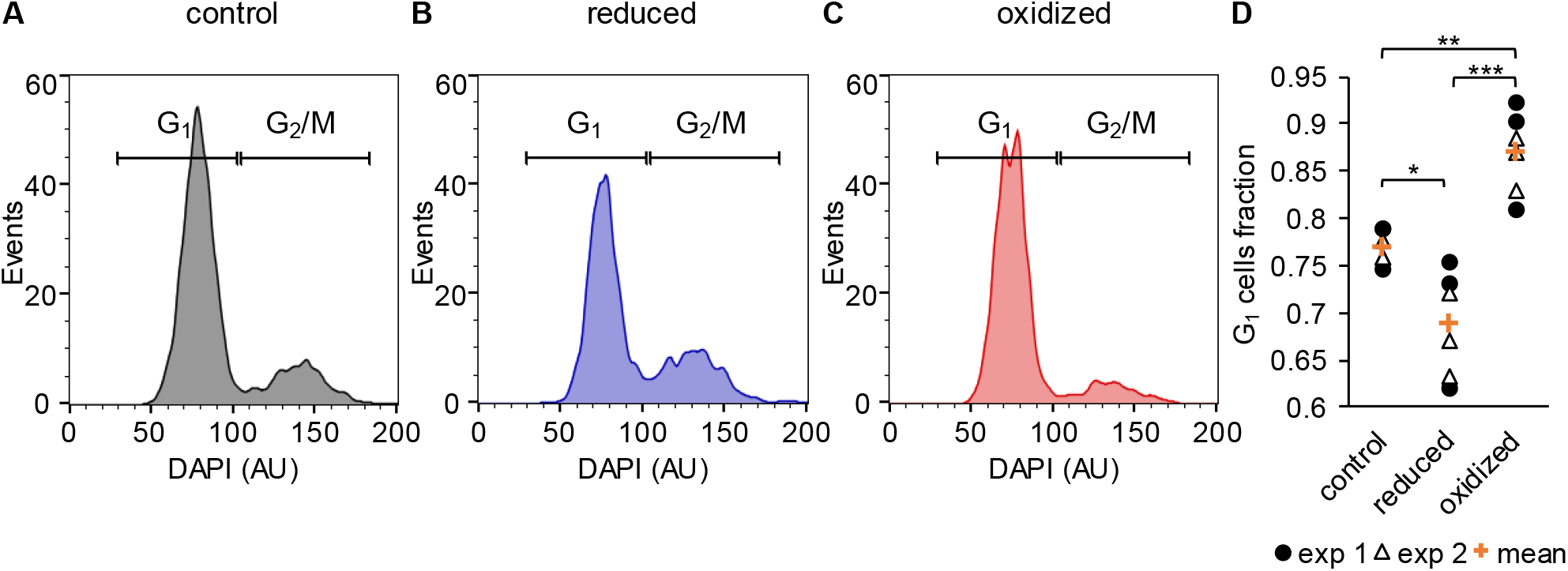
Cell cycle analysis of sorted reduced and oxidized subpopulations. (***A-C***) Cell cycle analysis of FACS sorted control untreated cells (A) and of “reduced” (B) and “oxidized” (C) subpopulations that were sorted 30 min post 80 μM H_2_O_2_ treatment, based on chl-roGFP OxD. DAPI staining was used for DNA content quantification, gates used to discriminate G_1_ (2 genome copies) and G_2_/M (4 copies) are marked. (***D***) Fraction of cells within the G_1_ gate (marked in A-C) in sorted subpopulations. Data points of n=6 biological repeats of 2 independent experiments (exp 1 and 2 marked with circles and triangles respectively) and the mean (orange plus sign) are shown, 1200-2000 cells analyzed per sample. *P*-values: *=0.011, **=0.0024, ***=0.0001. Tukey test was used for comparisons with control, paired t-test was used for comparing “reduced” and “oxidized” subpopulations.

### The fraction of “oxidized” cells is age dependent

To further explore the physiological factors that may affect the bi-stable distribution, we investigated the effects of different growth phases on oxidation of chl-roGFP in response to H_2_O_2_. Since aging has been associated with changes in redox states in several model systems^40–42^, we hypothesized that culture growth phase may affect the susceptibility of the population to oxidative stress. We challenged cultures originating from the same batch at different growth phases with 0-100 μM H_2_O_2_ after adjusting cell concentrations between growth phases (see methods, Fig. 6 A). Distinct “oxidized” and “reduced” subpopulations were detected in all growth phases examined, and in general older cultures had larger fractions of “oxidized” cells (Figures 6 B, S10 and S11). Following 80 μM H_2_O_2_ treatment, in early exponential culture 24.3 ± 0.8% of the cells were within the “oxidized” subpopulation, compared to the larger fraction in late exponential phase (43.6 ± 3.1%), while in stationary cultures the entire population was oxidized (*P*<0.001 for all comparisons, Tukey test, Fig. 6 B and Fig. S11 D). At a lower H_2_O_2_ concentration of 50 μM almost all exponential cells were within the “reduced” subpopulation, while most of the stationary cells were within the “oxidized” subpopulation, and more cells were oxidized in mid (85.9 ± 0.6%) compared to early stationary cultures (74.3 ± 2%, *P*<0.001, Tukey test, Fig. 6 B and Fig. S11 C). However, at stationary phase many cells were excluded from the analysis due to low roGFP fluorescence levels, which may lead to a bias in subpopulations proportions (Fig. S10, table S1).

**Fig. 6.**
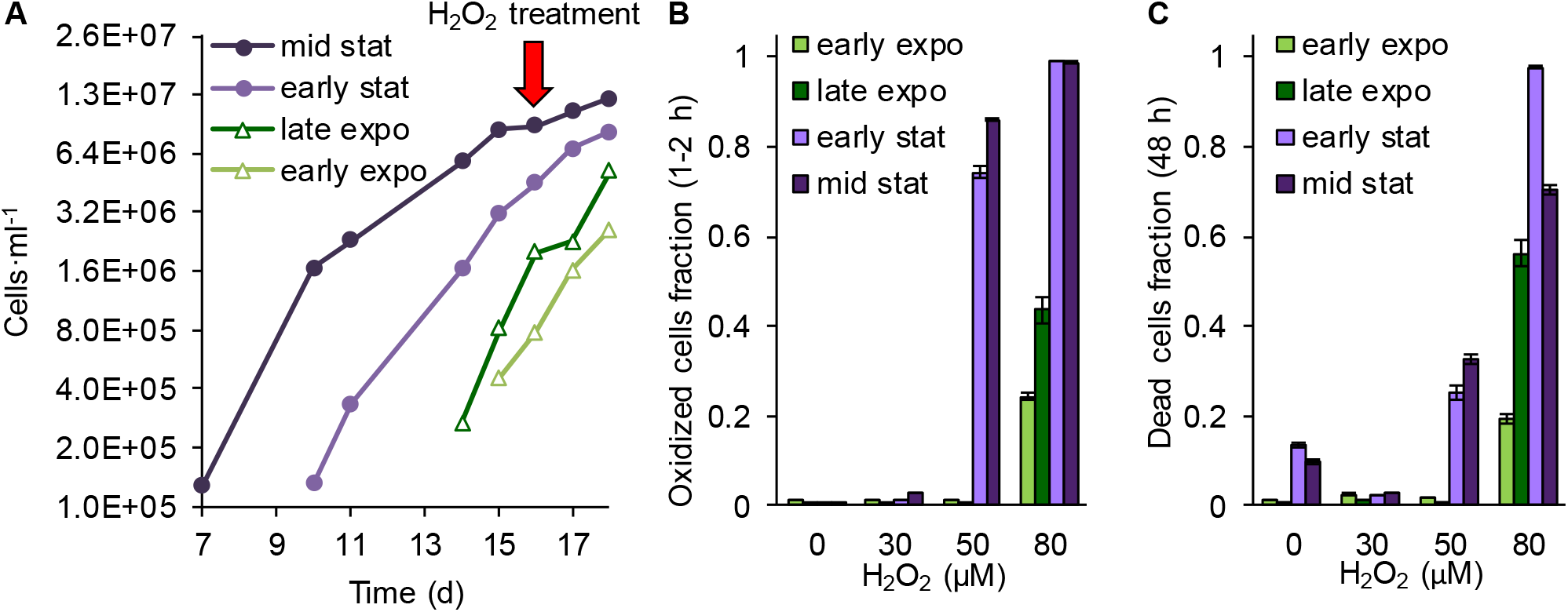
Culture growth phase affects chl-roGFP subpopulations proportions and H_2_O_2_ sensitivity. The effects of growth phase on chl-roGFP oxidation and mortality were measured using flow cytometry in early exponential (light green), late exponential (dark green), early stationary (light purple), and mid stationary cultures (dark purple). (***A***) Culture growth curve of chl-roGFP strain. The culture batch was sequentially diluted over time starting at day 0 to generate cultures at different growth phases, n=1 (see also Fig. S12 A). Red arrow marks the day of H_2_O_2_ treatment, cell concentration was adjusted prior to treatment. (***B***) The fraction of cells within the “oxidized” subpopulation 1 h post treatment with 0-80 μM H_2_O_2_. Mean ± SEM, n=3 biological repeats. (***C***) The fraction of dead cells 48 h post H_2_O_2_ treatment as measured by positive Sytox staining. Mean ± SEM, n=3 biological repeats.

Similarly to the “oxidized” subpopulation, the fraction of dead cells was both H_2_O_2_ dose-dependent and age-dependent, with higher fraction of dead cells in stationary compared to exponential cultures under the same treatment (*P*≤0.0011 for 50 and 80μM H_2_O_2_ treatments, Tukey test, Fig. 6 C). Interestingly, the induction of cell death was delayed in stationary cultures and was detected only 48 h post treatment compared to exponential cells, which showed significant cell death at 24 h post treatment (Fig. S12). Taken together, these results demonstrate that the proportions of the subpopulations and subsequent mortality depend on metabolic changes that occur during aging.

### The bi-stable chloroplast redox response is light dependent

Photosynthesis is the major source for reductive power as well as ROS in algal cells, and exposure to dark was shown to increase sensitivity to oxidative stress in another marine diatom^18^. Therefore, we hypothesized that light regime will affect the bi-stable pattern of chl-roGFP following oxidative stress, and investigated the effects of short exposure to darkness during daytime. Cells were treated with 0-100 μM H_2_O_2_ and were immediately moved to the dark for 90 minutes, after which they were moved back to the light (dark treated, Fig. 7 and Fig. S13). These cells were compared to cells that were kept in the light during this time (light treated). The transition to the dark caused an immediate oxidation of the basal chl-roGFP OxD (without H_2_O_2_ treatment), reaching a peak within 15 minutes (Fig. 7 A, D). Then, while still under dark, chl-roGFP gradually reduced while maintaining a continuous distribution (Fig. 7 A, D). Upon shifting back to the light, chl-roGFP reduced within 2 minutes back to its basal state prior to dark exposure (Fig. 7 A, D). The dark mediated oxidation was specific to the chloroplast and was not detected in the nucleus (Fig. S14), demonstrating the organelle specificity of these redox fluctuations.

**Fig. 7.**
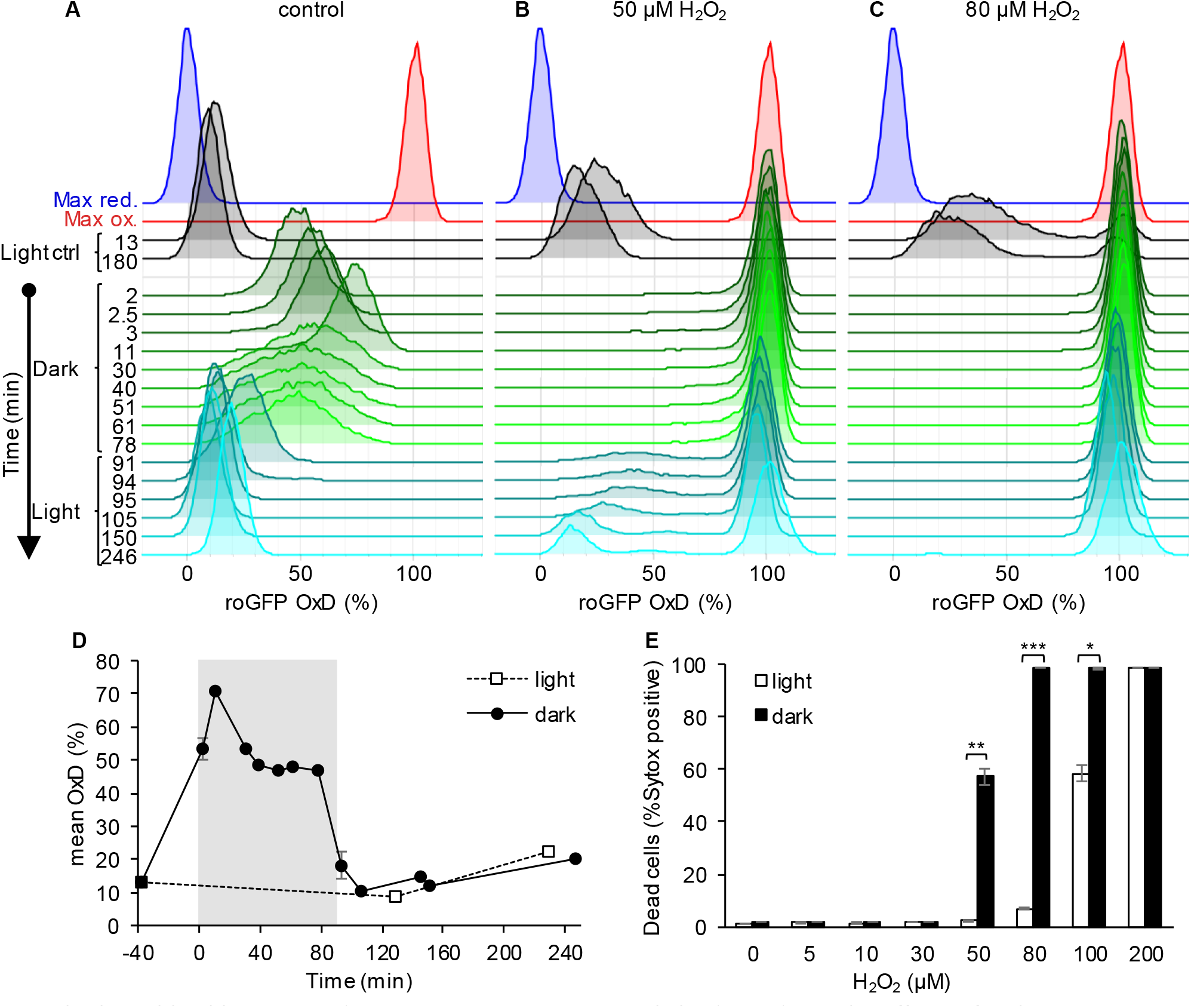
The bi-stable chl-roGFP oxidation in response to H_2_O_2_ is light-dependent. The effects of a short exposure to darkness during daytime on chl-roGFP oxidation patterns were examined. (***A-C***) Flow cytometry measurements of chl-roGFP OxD distribution in the population over time. Cells were treated with 0 μM (control, A), 50 μM (B), and 80 μM H_2_O_2_ (C), and were then transitioned to the dark at time 0 (within 5 minutes post H_2_O_2_ treatment). Cells were kept in the dark for 90 minutes (green) and were then transferred back to the light (cyan). The same H_2_O_2_ treatment without transition to the dark (light ctrl, black) and maximum oxidation (200 μM H_2_O_2_, red) and reduction (2 mM DTT, blue) are shown for reference. The experiment was done in triplicates that were highly similar, for visualization the first replica is shown. Each histogram consists of >8000 cells. (***D***) Mean ± SEM basal (control) chl-roGFP OxD over time of cells transitioned to the dark for 90 minutes (gray box) at time 0 (“dark”) and cells kept in light conditions (“light”), n=3 biological repeats. SEM lower than 0.5% are not shown. (***E***) Dead cells fraction 24 h post H_2_O_2_ treatment, with or without transition to the dark (“dark” and “light” respectively), as measured by positive Sytox staining. Data is shown as mean ± SEM, n=3 biological repeats. *P* values: *=0.0064, **=0.0026, ***=2·10^−5^, t-test.

The transition to the dark eliminated the bi-stable pattern of chl-roGFP oxidation in response to H_2_O_2_, and no distinct subpopulations were observed while cells were under darkness (Fig. 7 B-C and Fig. S13). The transition to the dark increased H_2_O_2_ sensitivity in the entire population, and following treatment of 80 μM H_2_O_2_ and transition to the dark chl-roGFP fully oxidized in the entire population and remained stably oxidized even after transition back to the light (Fig. 7 C). The bi-stable pattern was regained only upon transition back to the light, and only at lower doses of 30 μM and 50 μM H_2_O_2_, in which some or most cells were able to recover following this transition (Fig. 7 B and Fig. S13 C). In accordance with the higher chl-roGFP oxidation, “dark” treated cultures also exhibited higher mortality compared to “light” treated cells (*P*≤0.0064 for all pairs in 50-100μM H_2_O_2_ treatments, t-test, Fig. 7 E). Therefore, we conclude that the mechanism generating the bi-stable response in the chloroplast is light dependent and plays an important role in cell fate regulation in diatoms.

## Discussion

Our current understanding of the mechanisms that mediate acclimation to environmental stressors in marine microorganisms, including diatoms, is derived primarily from observations at the population level, neglecting any heterogeneity at the single-cell level. Averaging the phenotypes of a whole population can mask the co-existence of distinct subpopulations that employ diverse cellular strategies to improve the survival of this globally important phytoplankton group, as was shown in other microorganisms^43–45^. In this study, we established a novel system for studying phenotypic variability in the marine diatom *P. tricornutum* using flow cytometry and a microfluidics system for live imaging microscopy. Using organelle-specific measurements of E_GSH_ dynamics we assessed the *in vivo* redox state of individual diatom cells, and were able to detect a differential response to oxidative stress within the population. Based on this metabolic parameter, we identified two distinct subpopulations that emerged as an early response to oxidative stress, demonstrating the importance of phenotypic variability in cell fate regulation in diatoms.

We propose that in diatoms, the chloroplast E_GSH_ is involved in sensing specific environmental stress cues that induce oxidative stress, and in cell fate regulation (Fig. 8). Depending on the specific stress, ROS can accumulate at various sub-cellular compartments^16,17^, resulting in a shift of the population towards a more oxidized state. Cells that accumulate ROS above a certain threshold are likely to induce cell death with PCD-like hallmarks^16^, as observed in the “oxidized” subpopulation (Figs. 2 and 3). Cells that do not cross this threshold are able to recover and acclimate, as in the “reduced” subpopulation (Figs. 2 and 3). Using a microfluidics setup that allowed cell tracking throughout the entire dynamics during exposure to H_2_O_2_, we revealed such a “death threshold” based on early chl-roGFP oxidation (Fig. 3 G). This early response provided accurate cell fate predictions at the single-cell level. We propose that the balance between the prior metabolic state, antioxidant capacity, and the magnitude of the applied stress determines whether a cell will cross the “death threshold”, leading to a differential response within the population. Harsher stress conditions will have a stronger effect on the population, leading to more cells crossing the threshold and exhibiting early oxidation and subsequent cell death, as shown with increasing H_2_O_2_ doses (Fig. 1 M). Similarly, different basal metabolic states can increase the sensitivity of the entire population, leading to a larger fraction of “oxidized” cells even when they are exposed to lower H_2_O_2_ doses, as observed during culture aging (Fig. 6). The increased sensitivity in the dark (Fig. 7) further supports an involvement of the chloroplast in cell fate determination, suggesting that fluctuating light conditions experienced by diatoms in natural environments may greatly affect their susceptibility to additional stressors.

**Fig. 8.**
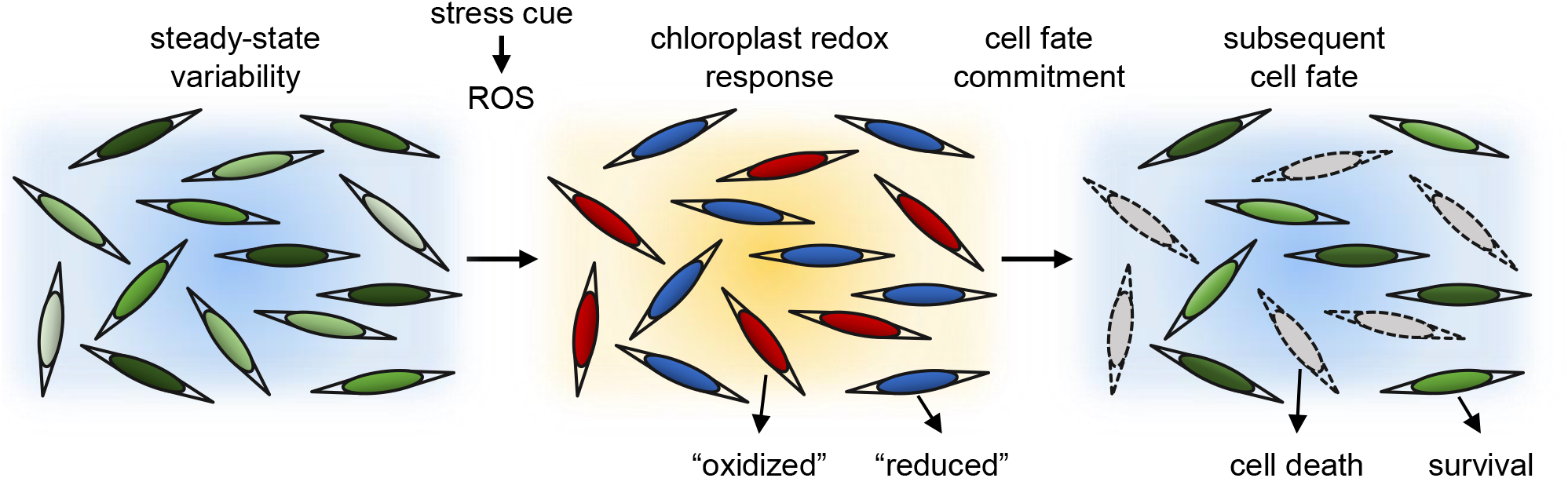
A conceptual model: phenotypic variability within diatom populations affects cell fate determination in response to stress conditions. At steady state conditions, cells within the population have diverse metabolic states due to various factors such as local ROS levels, antioxidant capacity, metabolic activity, growth phase and cell cycle position. We propose that this variability leads to a differential response to environmental stressors. Exposure to specific stress conditions leads to ROS accumulation at different subcellular compartments, including the chloroplast, which is used to sense the stress cue and regulate cell fate. Cells at a more susceptible metabolic state will accumulate high ROS levels and will subsequently die, as observed in the “oxidized” subpopulation. More resilient cells will exhibit milder oxidation and will be able to acclimate, as observed in the “reduced” subpopulation. Chloroplast E_GSH_ oxidation is an early stage in this stress response and precedes the commitment to cell fate.

Redox fluctuations in the chloroplast can serve as a rapid mechanism to perceive specific environmental cues, by regulating key metabolic pathways on the post-translational level, prior to gene expression. Analysis of the redox-sensitive proteome in *P. tricornutum* revealed over-representation of chloroplast-targeted proteins, that were also oxidized to a greater degree under H_2_O_2_ treatment as compared to other subcellular compartments^17,46^. Chloroplast E_GSH_ oxidation preceded the “point of no return” after which cell death was irreversibly activated (Fig. 2 D), supporting it has a role in sensing the stress cue. This “pre-commitment” phase provides an opportunity for cells to recover if conditions change during a narrow time frame of ∼30-100 min following oxidative stress (Fig. 2 D), before the cell has accumulated damage beyond repair or a PCD cascade was fully activated. This “pre-commitment” phase was shown previously in two diatom species, as exogenous application of the antioxidant GSH rescued diatom cells from otherwise lethal treatments of infochemicals or H_2_O_2_ only during the first hour^16,18^. These findings underscore the importance of redox regulation in chloroplast metabolic reactions, and its involvement in stress sensing and cell fate determination.

The role of the chloroplast in mediating PCD remains elusive, although mitochondria-generated ROS are known to play a key role in PCD in plants and animals^47,48^. This knowledge gap is even greater in unicellular marine algae, for which the molecular basis for the PCD machinery is largely unknown^8^. The chloroplast is a major source for generation of both ROS and reductive power to generate and recycle NADPH, thioredoxin and GSH^23^. Chloroplast-generated ROS were demonstrated to be involved in plants in retrograde signaling from the chloroplast and in hypersensitive response cell death^23,31,47,49^. A recent model suggested possible mitochondria-chloroplast cooperative interactions in the execution of ROS-mediated PCD^47^. In diatoms, mitochondrial ROS were linked to cell death in response to diatom-derived infochemicals^16^, and chloroplast E_GSH_ was shown to mediate changes in oxidative stress sensitivity upon light-dark transitions^18^. We propose that redox dynamics of both the mitochondria and the chloroplast are involved in cell fate regulation in diatoms.

The source of the single-cell variability observed in our system is yet to be further explored, but the results provide insights into factors that may drive it. Since clonal populations originating from single-cell isolates maintained the bi-stable chloroplast response, the variability does not result from genetic differences but rather from phenotypic plasticity (Fig. 4). It remains to be investigated whether the emergence of the subpopulations represents heterogeneity that occurs following exposure to stress, or rather a pre-existing variability within the population. Nevertheless, the difference in cell-cycle phase distribution between the subpopulations supports the latter. The combination of factors such as life history^33,50^, cell cycle phase^35,37^, cell age^51,52^, metabolic activity^53,54^, heterogeneous microenvironment^55^ and biological noise^43,56^ results in a distribution of different metabolic states within the population^57,58^. The metabolic state of the cell can affect its antioxidant capacity, therefore resulting in variability in sensitivity to oxidative stress as observed here. Importantly, the mechanism that generates variability in our system is light-dependent, as the bi-stable chl-roGFP pattern was abolished when the cells were under darkness and the entire population became more sensitive to oxidative stress (Fig. 7). The antioxidant capacity of a diatom cell depends on photosynthesis-generated NADPH, which is also used for GSH recycling. The transition to the dark may have compromised the biosynthesis and recycling of GSH, therefore enhancing sensitivity to oxidative stress^18^. Furthermore, stationary cultures were shown to have lower photosynthetic activity^59^, which results in a lower flux of NADPH generation and may lead to the increased sensitivity observed in stationary cultures (Fig. 6). Taken together, the source for heterogeneity could be variability in the flux of photosynthesis-derived reductive power.

Phenotypic variability can provide an important strategy to cope with fluctuating environments in microbial populations^57^. Future studies are required to investigate the possible tradeoff involved in maintaining high antioxidant capacity. For example, resilience to oxidative stress may come with a cost in the ability to sense environmental cues with high precision, as high ROS buffering capacity may mask milder ROS cues^46^. Co-existence of subpopulations with different susceptibilities to specific stressors can be viewed as a “bet-hedging” strategy of the population, enabling at least a portion of the population to survive unpredicted stress events and subsequently leads to a growth benefit at the population level^44,45,51,57^. A recent study demonstrated the benefit of phenotypic variability when NH ^+^ limitation increased the cell-to-cell variability in N_2_ fixation in the bacterium *Klebsiella oxytoca*, leading to improved growth under fluctuating conditions at the population level^44^. Phenotypic plasticity in response to stress was also shown in the toxic phytoplankton *Heterosigma akashiwo*, that improved its chances of evading deleterious turbulent conditions by diversifying its swimming directions^45^. Furthermore, phenotypic variability enables individual cells within isogenic populations to exhibit various cell fates, including PCD, despite possible detrimental effects at the single-cell level. In diatoms, phenotypic variability in cell size, shape and susceptibility to stress conditions were suggested^1,16,18,60^, but until now the experimental setups were not designed to study individuality in stress response. Redox-based phenotypic variability may provide a rapid and adjustable strategy to cope with unpredicted stress conditions as compared to relying only on genetic diversity.

The novel approaches developed here provide new insights into individuality in marine microbes, and enable studying dynamic processes at the single-cell level in diatoms and possibly other ecologically relevant microorganisms. The ecological importance of variability in the chloroplast redox state and the mechanisms that underlie differential sensitivity to oxidative stress are yet to be explored.

## Methods

### Culture growth

*P. tricornutum* accession Pt1 8.6 (CCMP2561 in the Provasoli-Guillard National Center for Culture of Marine Phytoplankton) was purchased from the National Center of Marine Algae and Microbiota (NCMA, formerly known as CCMP). Cultures were grown in filtered sea water (FSW) supplemented with F/2 media^61^ at 18°C with 16:8 hours light:dark cycle and 80 μmol photons m^−2^ sec^−1^ light intensity supplied by cool-white LED lights (Edison, New Taipei, Taiwan). Strains expressing roGFP were obtained as described previously^16,17^. Cultures were kept in exponential phase, experiments were performed in ∼0.5-1·10^6^ cells·ml^−1^.

### roGFP measurements

roGFP oxidation was measured over time following the addition of H_2_O_2_ or in untreated control using the ratio between two fluorescence channels, i405 and i488, by fluorescence microscopy (described below) and by flow cytometry using BD LSRII analyzer, BD FACSAria II and BD FACSAria III. The roGFP ratio (i405/i488) increases upon oxidation of the probe^62^. The oxidation degree of roGFP (OxD) was calculated according to Schwarzländer *et al.*^62^:

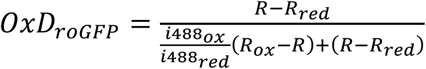

Where R is the roGFP ratio of i405/i488, Rred is the ratio of fully reduced form (15-50 minutes post treatment with 2 mM Dithiothreitol, DTT), R_ox_ is the ratio of the fully oxidized form (7-30 minutes post treatment with 200 μM H_2_O_2_), and i488_ox_ and i488_red_ are the i488 of the maximum oxidized and maximum reduced forms respectively. For sorting purposes, roGFP ratio was used, as exact OxD cannot be calculated prior to sorting, both parameters give similar partition between the subpopulations (data not shown). In flow cytometry measurements, i405 was measured using excitation (ex) 407 nm, emission (em) 530/30 nm or 525/25 nm, and i488 was measured using ex 488 nm, em 530/30 nm. Relative expression level of roGFP was measured by multiplication of i405 and i488, and was used to gate roGFP+ cells during analysis (Fig. S2 D). In sorting experiments, i405 and i488 were used instead to gate roGFP+ cells since relative roGFP expression could be calculated only post acquisition. Dynamic range of roGFP was calculated by ratio of Rox/Rred (tables S1, S2). Flow cytometry measurements were done under ambient light and temperature conditions, except for dark treatment during which samples were covered with aluminum foil. At time 0, H_2_O_2_ treatment was added from a freshly prepared 20 mM stock to *P. tricornutum* cultures to a final concentration of 5-200 μM. There were differences in roGFP fluorescence levels between the strains (Fig. S2 D). The mit-roGFP strain was less informative due to the lower roGFP fluorescence and therefore lower signal to noise ratio (SNR) and smaller dynamic range (Figs. S2 D and S4, table S2). It is important to note that the average leakage of chlorophyll auto-fluorescence (AF), especially into the i405 channel, increased over time starting ∼116 minutes post 80 μM and 100 μM H_2_O_2_ treatments in all strains, but was most prominent in the mit-roGFP strain (Figs. S16 and S17). AF leakage was not linearly correlated with AF in the chlorophyll channel, and therefore could not be corrected at the single cell level. It may introduce a bias towards oxidation in roGFP oxidation calculations at later time points and can explain values above 100% OxD, but at early time points these effects were negligible at least in the nuc-roGFP and chl-roGFP strains (Figs. S16 and S17).

### Cell death analysis

For cell death analysis, samples were stained with a final concentration of 1 μM Sytox Green (Invitrogen), incubated in the dark for 30-60 min at RT and analyzed using an Eclipse flow cytometer (ex 488 nm, em 525/50 nm). Unstained samples were used as control to discriminate background signal. In microfluidics experiments, Sytox was dissolved in fresh media (FSW+F/2) to a final concentration of 1-2 μM, and inserted into the system at 21.5-23 hours post treatment, without changing the flow rate (1 μl/min). Fresh stain was continuously flowing through the system for at least 1.5 hours during which cells were imaged for Sytox staining (ex 470/40 nm, em 525/50 nm) as described below. For Sytox staining analysis, images of 30-106 minutes incubation time were used based on highest staining and best focus.

### Sorting of subpopulations and generation of clonal populations

To measure survival and generate clonal populations originating from different subpopulations, cells expressing chl-roGFP of the “oxidized” and “reduced” subpopulations were sorted according to their roGFP ratio at different times post 80 μM H_2_O_2_ treatment using BD FACS AriaII and BD FACS AriaIII. The “oxidized” and “reduced” subpopulations gates were based on visible separation and avoiding cells near intermediate values (Fig. 2 C). Untreated control cells were sorted according to positive roGFP fluorescence, the gate upstream of the subpopulations gates. To avoid oxidation due to darkness within the FACS machine, sorting times were minimized and the “oxidized” subpopulation was sorted first followed immediately by sorting of the “reduced” subpopulation. However, in longer sorting sessions as for the Sytox and cell cycle analyses, some cells within the sorted “oxidized” subpopulation may have been oxidized due to the combined effect of H_2_O_2_ and exposure to the dark within the FACS. For Sytox analysis of cell death post sorting, 10,000 cells/well were sorted into fresh media (FSW+F/2) in triplicate biological repeats. For single cell survival 1 cell/well was sorted into 96-well plates containing either “agar” (1.5% agarose + FSW/2 + F/2 + antibiotics) or “liquid” (FSW+F/20) fresh media. Cells grown in liquid were further diluted and then spotted on agar plates. 5-9 weeks post sorting the plates were scanned and colonies were counted manually, assuming each colony originates from a single surviving cell. Since survival was highly similar in liquid and in agar the results of these two methods were combined together. Each method was done in biological triplicates per medium type per experiment, data is shown for two independent experiments for time points 30 and 100 min and one experiment for the 60 min time point. For generation of clonal populations, single cells sorted into liquid medium were used. Clones were exposed to 80 μM and 100 μM H_2_O_2_ ∼3-6 weeks post sorting, and their chl-roGFP oxidation was measured using flow cytometry. A total of 18 “control”, 29 “oxidized” and 32 “reduced” clones were examined in two independent experiments.

### Microfluidics chip preparation

Microfluidics chip design was based on Shapiro *et al.*^63^, and was modified for *P. tricornutum* cells. Each chip contained 4 channels of 2 cm length X 0.2 cm width X 150 μM height with one circular widening of 0.4 cm diameter, with a total volume of ∼12.7 μl per channel (Fig. 3 H and Fig. S6 B). The microfluidics chip was etched into a silicone elastomer (Sylgard 184, Dow Corning) using soft lithography. Silicone elastomers were prepared by mixing the two components in a 10:1 ratio and were poured onto the dust-free wafer, de-aired in a desiccator to eliminate air bubbles, and incubated overnight at 60 °C for curing to generate the Polydimethylsiloxane (PDMS) microfluidics chips. Inlet and outlet holes were punched at both ends of each channel using a 1 mm biopsy punch (AcuDerm, FL, USA). The PDMS chip was placed on the clean surface of a new glass microscope 60×24 mm cover slip using plasma bonding with a BD-20AC corona treater (Electro-Technic Products) followed by heating of 100°C for >15 minutes to ensure covalent bonding of the PDMS and the glass.

### Microfluidics live imaging

Light and epifluorescence microscopy imaging was performed using a fully motorized Olympus IX81 microscope (Olympus) equipped with ZDC component for focus drift compensation, using 20X air objective (numerical aperture 0.5) and Lumen 200PRO illumination system (Prior Scientific). Images were captured using a Coolsnap HQ2 CCD camera (Photometrics, Tuscon, AZ, USA). The microfluidics chip was mounted on a motorized XY stage (Prior Scientific, MA, USA) with a temperature-controlled inset (LCI, Korea) set to 18°C (Fig. S6 C). The outlet tubes were connected to syringe pumps (New Era Pump Systems, NY, USA) set to withdraw mode, using negative pressure for flow generation. The inlet tubes were connected to Eppendorf reservoirs, containing the fluid to be inserted into the system. Experiment layout is shown in Fig. S6 A. Chambers were washed with at least 500 μl of pure ethanol, then double-distilled water and then fresh media prior to the introduction of cells. Cells were introduced into the system and settled on the glass bottom. Flow rate was kept at 1 μl/min for the duration of the experiment, except during cell introduction (100 μl/min), cell settlement (up to 20 μl/min with occasional stops), and treatment introduction (10 μl/min for the initial 10 minutes for rapid replacement of media). Following settlement and at least 1 hour after cells were introduced to the system, cells were imaged for roGFP measurements (roGFP i405: ex 405/20 nm, em 525/50 nm; roGFP i488: ex 470/40 nm, em 525/50 nm), chlorophyll auto-fluorescence (ex 470/40 nm; em 590 lp) and bright field (BF, without a condenser). Each chip contained 4 chambers that were imaged sequentially: chl-roGFP control, chl-roGFP 80 μM H_2_O_2_ treated, WT 80 μM H_2_O_2_ treated and WT control (WT strain can be used to monitor auto-fluorescence changes and leakage during experiments). In each chamber, 5-6 different fields were imaged every 20 minutes over the course of >24 hours to avoid photo-toxicity. Ambient light was provided during light period using the microscope’s BF illumination without a condenser, light intensity ranging between 34 (at the very edge, outside the imaging region) to 80 (center) μmol photons m^−2^ sec^−1^. No images were obtained during the night to avoid disturbance to the diurnal cycle. After imaging the basal state of the cells, treatments of either 80 μM H_2_O_2_ dissolved in fresh media (FSW+F/2) or fresh media control were introduced to the system continuously for ∼2.5-3 hours, after which they were gradually washed away by fresh media. To quantify cell death, Sytox green was introduced into the system at 21.5-23 hours post treatment (see above) and was imaged using the roGFP i488 channel with shorter exposure time. The Sytox signal was stronger than the roGFP i488 and was localized mainly to the nucleus, enabling a clear separation between the two signals (Fig. 3 B, D). Only a small fraction of cells within the control treatment were Sytox positive (0.0054%), indicating that cells remained viable in this experimental setup. Furthermore, cells of the “reduced” subpopulation and of control treatment were able to proliferate, further demonstrating their viability under these conditions (movies S1, S2).

### Image analysis

Image analysis was performed using a designated MATLAB based script (see overview in Fig. S15) that is available on GitHub: https://github.com/aviamiz/ITRIA. Images were imported using bio-formats^64^. Then, image registration for XY drift correction was done using the Image Stabilizer plugin^65^ for FIJI (Fiji Is Just ImageJ) and using MIJI^66^ to access FIJI from MATLAB. Then images were normalized by bit-depth. Background subtraction was done based on mean value of a user-defined region of interest (ROI) that did not include cells. All fluorescence channels (i405, i488 and chlorophyll) were thresholded by a user-defined value to generate masks of positive expression. The roGFP relative expression level was calculated pixel-by-pixel by multiplication of the i405 and i488, only at pixels that were co-localized in the i405 and i488 masks. Then, roGFP relative expression (i405 * i488) was thresholded in order to include only pixels with high enough signal, based on a user-defined threshold. The roGFP ratio and OxD were calculated pixel-by-pixel as described above, pixels that were not included in the roGFP expression mask were excluded and set to NaN (not a number). For values of maximum oxidation and reduction of roGFP, cells were imaged in the same microfluidics imaging setup following treatments of 200 μM H_2_O_2_ and 2 mM DTT respectively (see “roGFP calculations”). Cell segmentation was based on i405 (chl-roGFP strain) or chlorophyll (WT strain) masks and fluorescence intensity using watershed transformation. Cells were filtered based on area, major and minor axis length, and eccentricity in order to exclude clumps of cells and doublets. Cell tracking was adapted and modified from a MATLAB code kindly provided by Vicente I. Fernandez and Roman Stocker^67,68^. In short, particles were tracked based on minimizing the distance between particle centroids in adjacent frames within a distance limit. Sytox analysis was based on a user defined threshold and co-localization of the Sytox with the extended cell region within the cell segmentation mask. Images from the same experiment were analyzed using the same values for all thresholds and parameters, except for Sytox analysis in which the threshold was adjusted manually to validate correct assignment of cell-fate and to avoid effects of focus differences. Cells that were not detected in the frame used for Sytox analysis or were not tracked for at least 6 consecutive frames were excluded from further analysis. The 74% OxD threshold used for early discrimination between the subpopulations and for cell fate prediction (Fig. 3 E-G and Fig. S7) was based on logistic regression modelling of cell fate at the end of the experiments as a function of chl-roGFP OxD 40 min post 80 μM H_2_O_2_ treatment (Fig. S8). Only a minor fraction of the cells exhibited intermediate roGFP OxD of 60-80% (10 tracks out of 754, 1.33%) or seemed to recover and shifted from the oxidized subpopulation to the reduced subpopulation overnight (3 tracks, 0.4%, Fig. 3 F). However, these rare cases require manual validation and may result from improper segmentation of adjacent cells with different oxidation states or due to tracking mistakes. The observed roGFP OxD of more than 100% oxidation in some cells could result from increased auto-fluorescence leakage to the i405 channel at later times post treatment (see Figs. S16 and S17).

### Cell cycle analysis

Cell cycle analysis was based on Huysman *et al.*^39^ and modified for sorted cells. 30,000 cells of “oxidized” and “reduced” sub-populations were sorted 30 min post 80 μM H_2_O_2_ treatment into 260 μl 80% ethanol kept at 5°C, reaching a final concentration of 70% ethanol. Control untreated cells and synchronized cells (20 h dark, as previously described^39^) were sorted based on positive roGFP fluorescence. Cells were then gently mixed and kept at 4°C until further processing. Then, 500 μl of 0.1% bovine serum albumin in phosphate-buffered saline (PBS) was added to improve pellet yield, and samples were centrifuged at 4000 rcf at 4°C for 10 min to discard supernatant. Cells were then washed with PBS, re-suspended and stained with 4’,6-diamidino-2-phenylindole (DAPI, Sigma) at a final concentration of 10 ng/ml. Samples were analyzed using BD LSRII analyzer, with ex 355 nm and em 450/50 nm. Synchronized cells were used as a reference to validate the gates for G_1_ and G_2_/M phases (data not shown). S phase was not clearly detected in this analysis, and therefore both gates likely include also S phase cells.

### Growth phase

The same culture batches of WT and chl-roGFP strains were diluted sequentially starting at day 0, and cell concentration was measured using Multisizer™ 4 COULTER COUNTER (Beckman Coulter) (Fig. 6 A and Fig. S12 A). On day 16, cultures at 4 different growth phases (early exponential, late exponential, early stationary and mid stationary, Fig. 6 A) were treated with 0-80 μM H_2_O_2_ and their chl-roGFP oxidation response and subsequent mortality were measured as described above. To exclude effects of different cell concentrations, 1-2 h before treatment cell concentrations were adjusted to 6.7·10^5^ cells·ml^−1^ (concentration of early exponential culture) by centrifugation and resuspension in fresh media. In stationary cultures a small subpopulation (2-3%) with high AF was observed using ex 405 nm em 450/50 nm and exhibited OxD values >110%, and was therefore excluded from the analysis (table S1).

### Statistics

All statistical analyses were done in R. ANOVA was used for multiple comparisons, and then Dunnett test or Tukey test were performed were applicable. For comparisons of two samples, t-test was used, and paired t-test was used where applicable. Values are represented as mean ± SEM unless specified otherwise. Box-plot was generated using the web tool BoxPlotR http://shiny.chemgrid.org/boxplotr/^69^ using Tukey whiskers, which extend to data points that are less than 1.5 x Interquartile range away from 1st/3rd quartile.

### Data availability

All relevant data supporting the findings of the study are available in this article and its Supplementary Information, or from the corresponding author upon request. Custom-made code for image analysis is available from the corresponding author upon request.

## Author contributions

A.M. and A.V. designed research; A.M. performed all experiments; A.M. and O.H.S. designed the microfluidics setup; A.M. developed methods and code for image analysis; A.M, S.G.v.C., S.R. and A.V. analyzed data; and A.M. and A.V. wrote the manuscript with help from all the authors.

## Acknowledgments

We thank Dr. Ron Rotkopf from the Bioinformatics Unit at the Life Sciences Core Facilities, Weizmann Institute of Science for assisting with the statistical analysis. We thank Dr. Vicente I. Fernandez and Prof. Roman Stocker for help with the cell tracking algorithm. We thank Dr. Uri Sheyn and the flow cytometry unit at the Life Sciences Core Facilities, Weizmann Institute of Science for technical help with FACS operation. We thank Jenny Mizrahi for proofing the manuscript. This research was supported by the Israeli Science Foundation (ISF) (grant #712233) awarded to AV.

## Supplementary Information

**Fig. S1.**
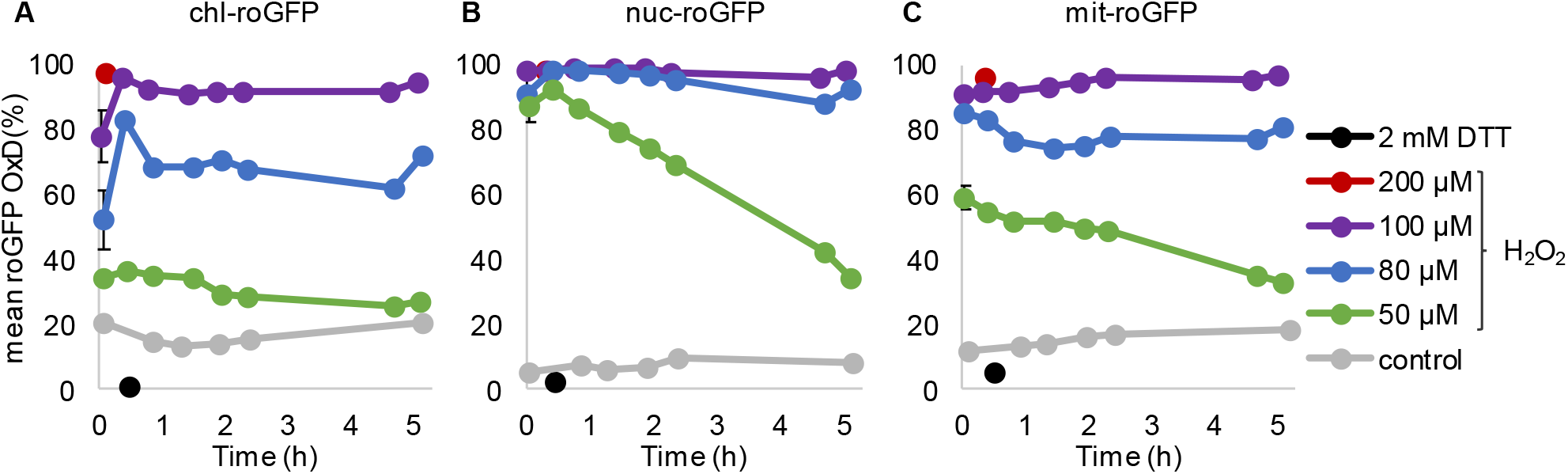
Organelle specific roGFP mean oxidation in response to H_2_O_2_ application in *P. tricornutum*. Flow cytometry measurements of the mean ± SEM roGFP OxD in the population over time in response to oxidative stress in chl-roGFP (A), nuc-roGFP (B) and mit-roGFP (C) *P. tricornutum* strains, n=3 biological repeats. Oxidation dynamics were measured following treatments with different H_2_O_2_ concentrations: 50 μM (green), 80 μM (blue), 100 μM (purple) and without treatment (control, gray). Maximum reduction (2 mM DTT, black) and maximum oxidation (200 μM H_2_O_2_, red) are shown for reference. Error bars below 1.5% are not shown.

**Fig. S2.**
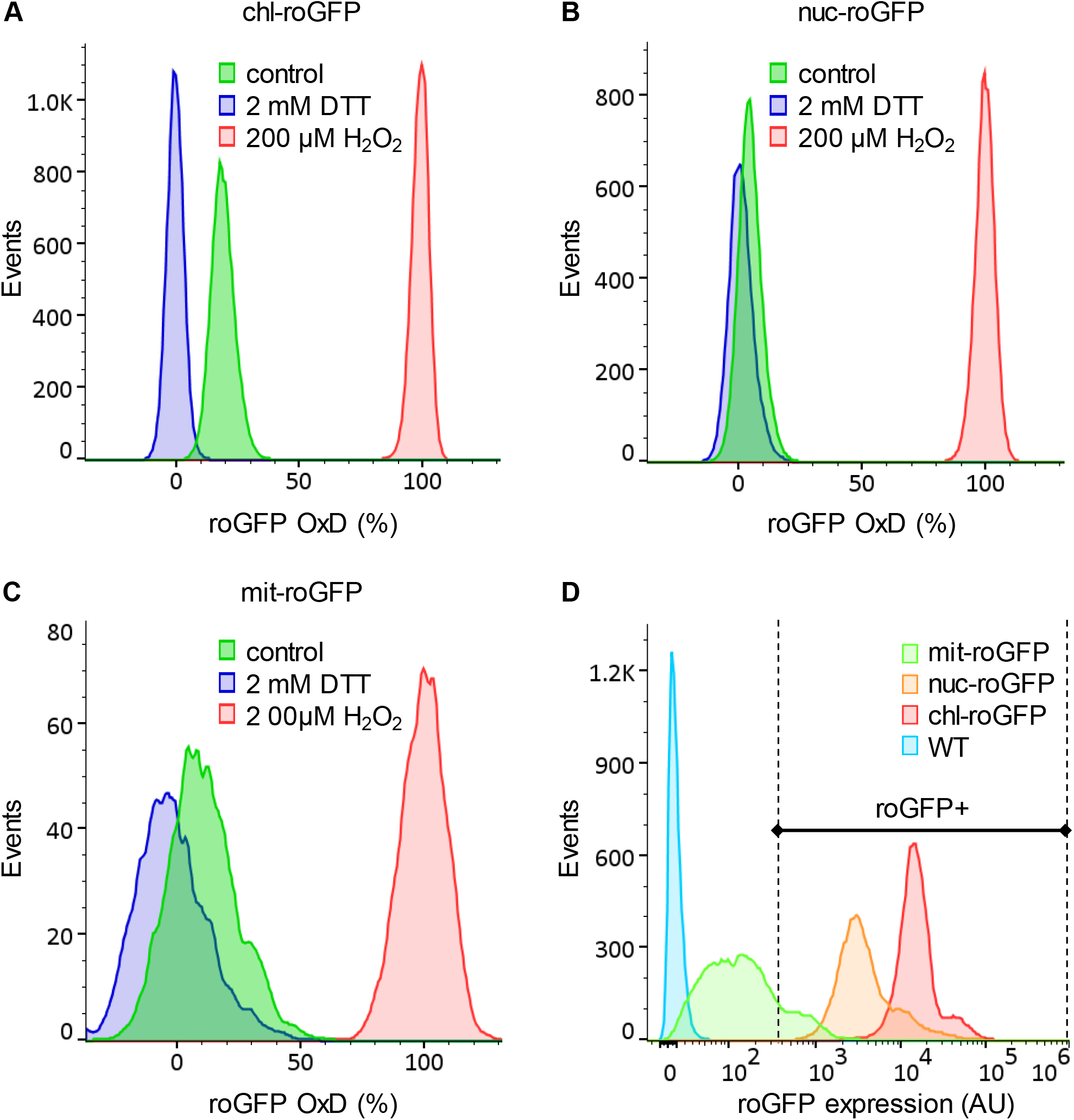
Distribution of roGFP oxidation in different subcellular compartments of *P. tricornutum* cells at steady state. Flow cytometry measurements of roGFP OxD and expression levels in *P. tricornutum* cells expressing roGFP targeted to the chloroplast (chl-roGFP, A), nucleus (nuc-roGFP, B) and mitochondria (mit-roGFP, C). (***A-C***) The distribution of roGFP OxD in the population at steady state conditions (without treatment, green) and following treatments of maximum oxidation (200 μM H_2_O_2_, red) and maximum reduction (2 mM DTT, blue). (***D***) Relative roGFP expression levels (AU) based on fluorescence intensity in chl-roGFP (red), nuc-roGFP (orange), mit-roGFP (green) and WT (no roGFP, cyan) strains. Relative roGFP expression was calculated by multiplication of the i405 and i488 (see methods). A threshold was set to exclude cells with low roGFP expression from roGFP oxidation calculations, shown as a gate with the label “roGFP+”. Each histogram consists of >12,000 cells (A, B, D) or >2,400 cells (C, due to lower roGFP expression). The experiment was done in triplicates, for visualization purposes one representative repeat is shown.

**Fig. S3.**
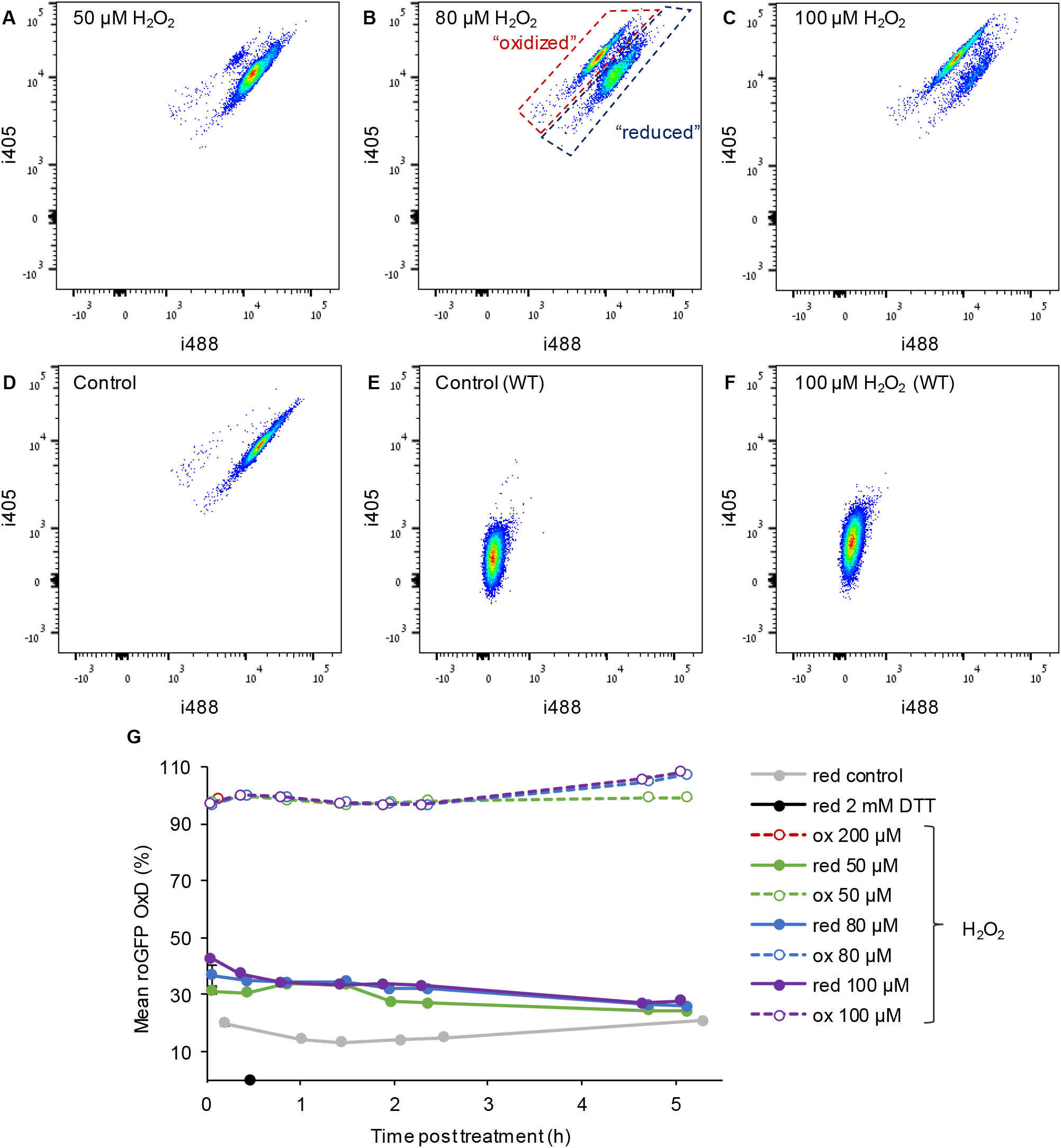
Oxidation of chl-roGFP reveals distinct subpopulations in response to H_2_O_2_ treatment. (***A-F***) Density plots of flow cytometry fluorescence measurements of i405 over i488 (arbitrary units, see methods) of chl-roGFP (A-D, roGFP positive cells) and WT cells that do not express roGFP (E-F, chlorophyll positive cells) 83-88 minutes post H_2_O_2_ treatments of 50 μM (A), 80 μM (B), 100 μM (C, F) and untreated control (D, E). The ratio i405/i488 represents the roGFP oxidation state, and it increases upon oxidation. The “oxidized” and “reduced” chl-roGFP subpopulations are marked in red and blue respectively in B. (***G***) Flow cytometry measurements of roGFP OxD of the “oxidized” (dashed, empty symbols) and “reduced” (solid, full symbols) subpopulations over time post 0-200 μM H_2_O_2_ treatments. Maximum reduction following 2 mM DTT (black) is shown for reference. Results are shown as mean ± SEM, n=3 biological repeats.

**Fig. S4.**
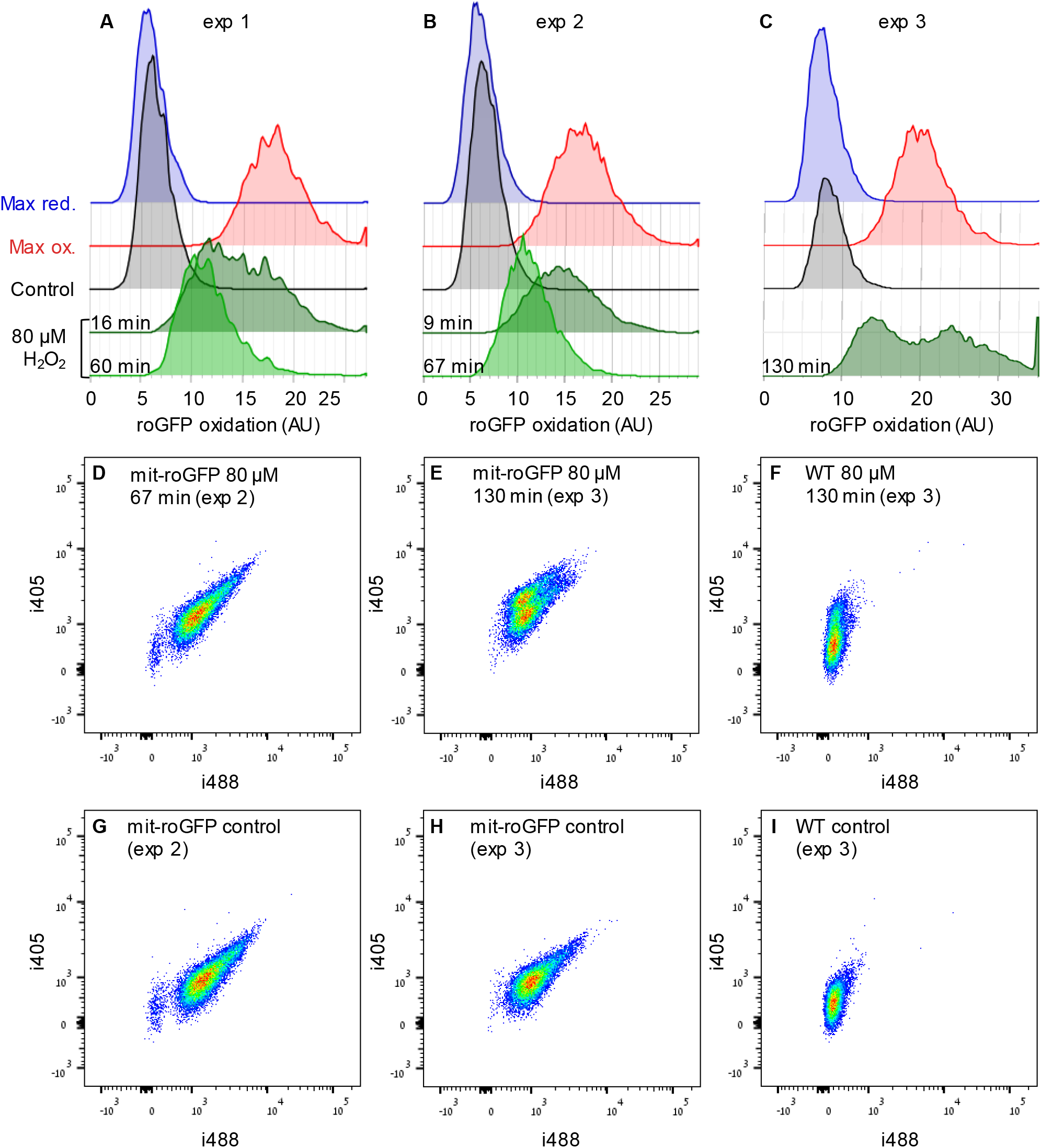
Heterogeneity in mit-roGFP response to H_2_O_2_. Flow cytometry measurements of mit-roGFP oxidation in response to 80 μM H_2_O_2_ in 3 independent experiments (in addition to the experiment shown in Fig. 1): exp 1 (A), exp 2 (B, D, G) and exp 3 (C, E, F, H, I). (***A-C***) Distribution of mit-roGFP OxD at different times post 80 μM H_2_O_2_ (green). Measurements of untreated control (black) and following maximum oxidation (red, 200 μM H_2_O_2_) and maximum reduction (blue, 2 mM DTT) are shown for reference. (***D-I***) Density plots of roGFP fluorescence measurements of i405 vs. i488 of mit-roGFP (D, E, G, H) and WT (F, I) strains at different times post 80 μM H_2_O_2_ treatment (D-F) or untreated control (G-I). The ratio i405/i488 represents roGFP oxidation state (see methods). WT is shown for auto-fluorescence (AF) leakage reference, demonstrating the effect of lower expression level in mit-roGFP strain. A differential response was observed in exp 3 and in the exp shown in Fig. 1 K, L, but in exp 1 and 2 no distinct subpopulations were observed. This differential response was clearly observed only at later time points, which were not measured in exp 1 and 2, and may result from auto-fluorescecne leakage (see Fig. S16 and S17). Measurements were done in triplicates, one repeat is shown for visualization.

**Fig. S5.**
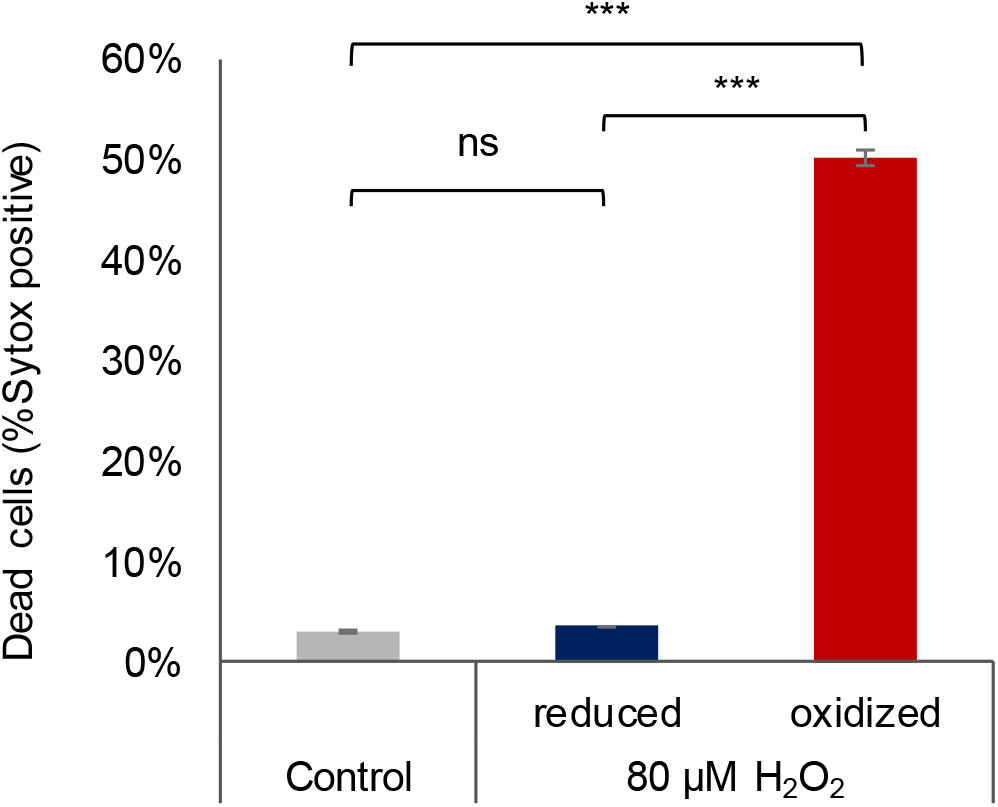
Mortality of sorted subpopulations following H_2_O_2_ treatment. Cell death was measured using Sytox staining 24 h post sorting of enriched “oxidized” and “reduced” subpopulations or untreated control (see methods). Subpopulations were sorted 112 min post 80 μM H_2_O_2_ treatment. Data is shown as mean ± SEM, n=3 biological repeats, each consists of >3700 cells. (***) *P*<0.001; (ns) non-significant (*P*=0.73), Tukey test.

**Fig. S6.**
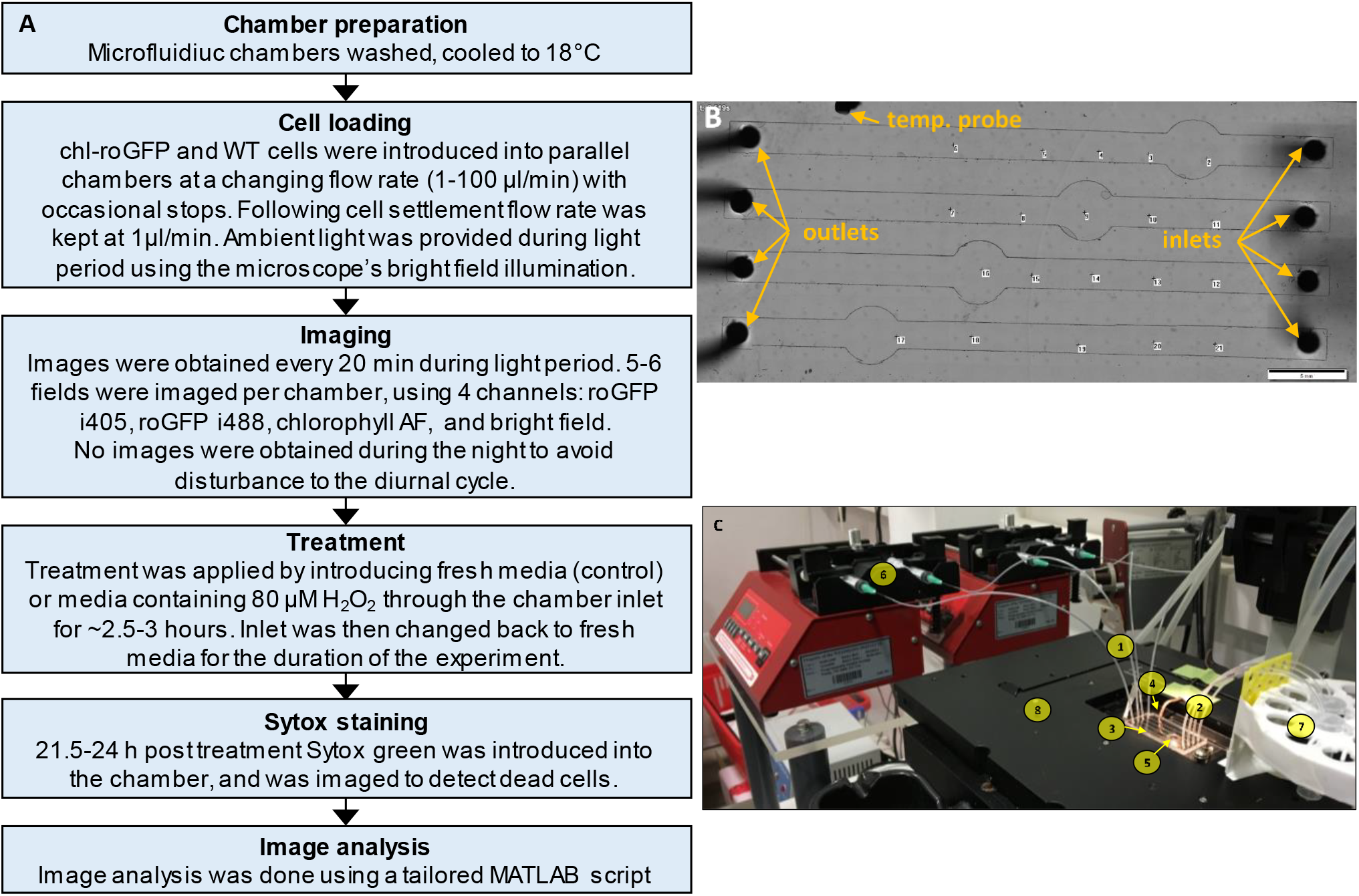
Microfluidics experimental layout. (***A***) Schematic representation of microfluidics experiments. (***B***) Bright field image of the microfluidics chip (4x magnification) with markings of example imaging positions (dots), temperature probe, inlets and outlets. Scale bar = 5 mm. (***C***) The microfluidics experimental setup. 1: outlet tubes; 2: inlet tubes; 3: microfluidics chamber; 4: temperature electrode; 5: microscope objective; 6: syringe pumps; 7: inlet reservoirs; 8: motorized stage with controlled temperature.

**Fig. S7.**
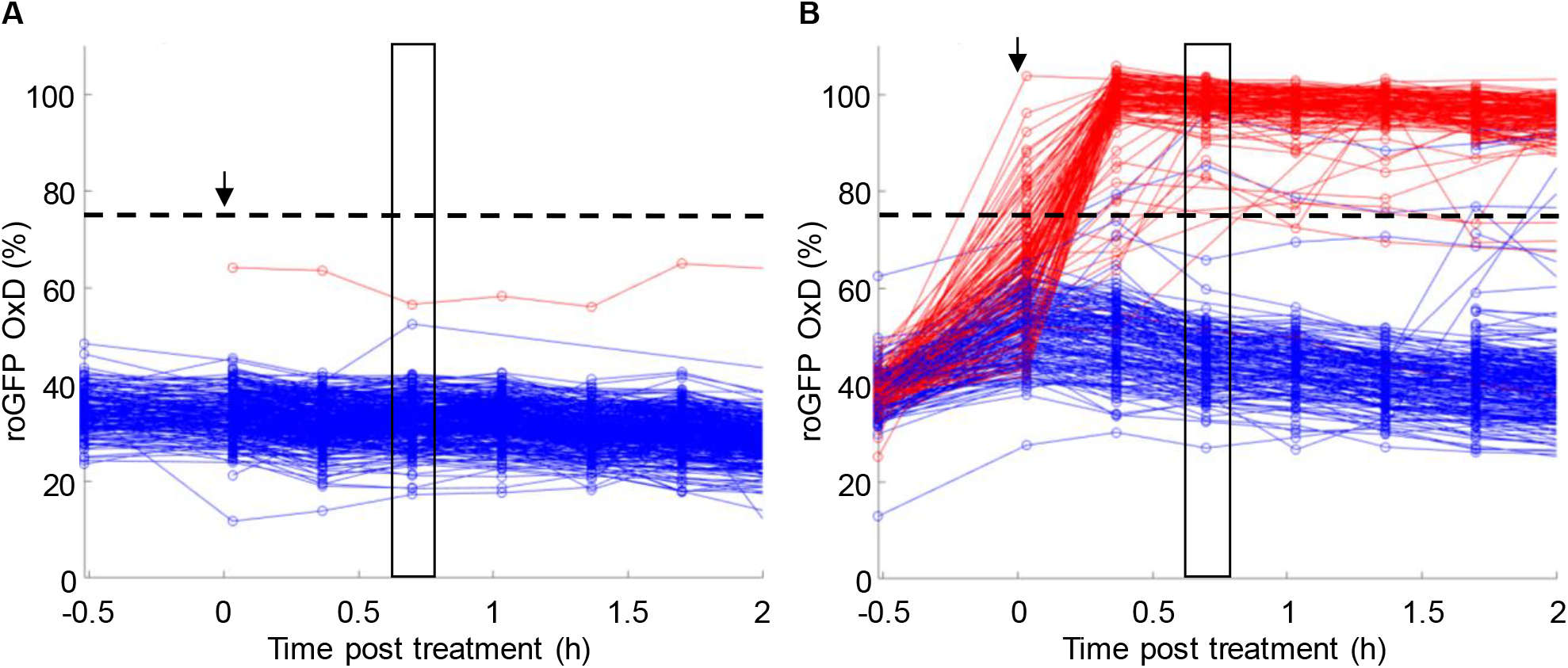
Tracking early chl-roGFP oxidation dynamics in single cells following oxidative stress using *in vivo* imaging in a microfluidics setup. OxD of chl-roGFP per cell over time following treatment with either fresh media (control; A) or 80 μM H_2_O_2_ (B) of cells imaged over time using microfluidics and epifluorescence microscopy. Treatment was introduced at time 0 (arrow). Color is based on subsequent cell fate as measured by Sytox staining ∼23 h post treatment: blue – alive, red – dead. N≥250 cells from at least 3 different fields. Horizontal dashed line represents the threshold used for cell fate predictions. The black box marks the time point used for cell fate prediction analysis in Fig. 5 G.

**Fig. S8.**
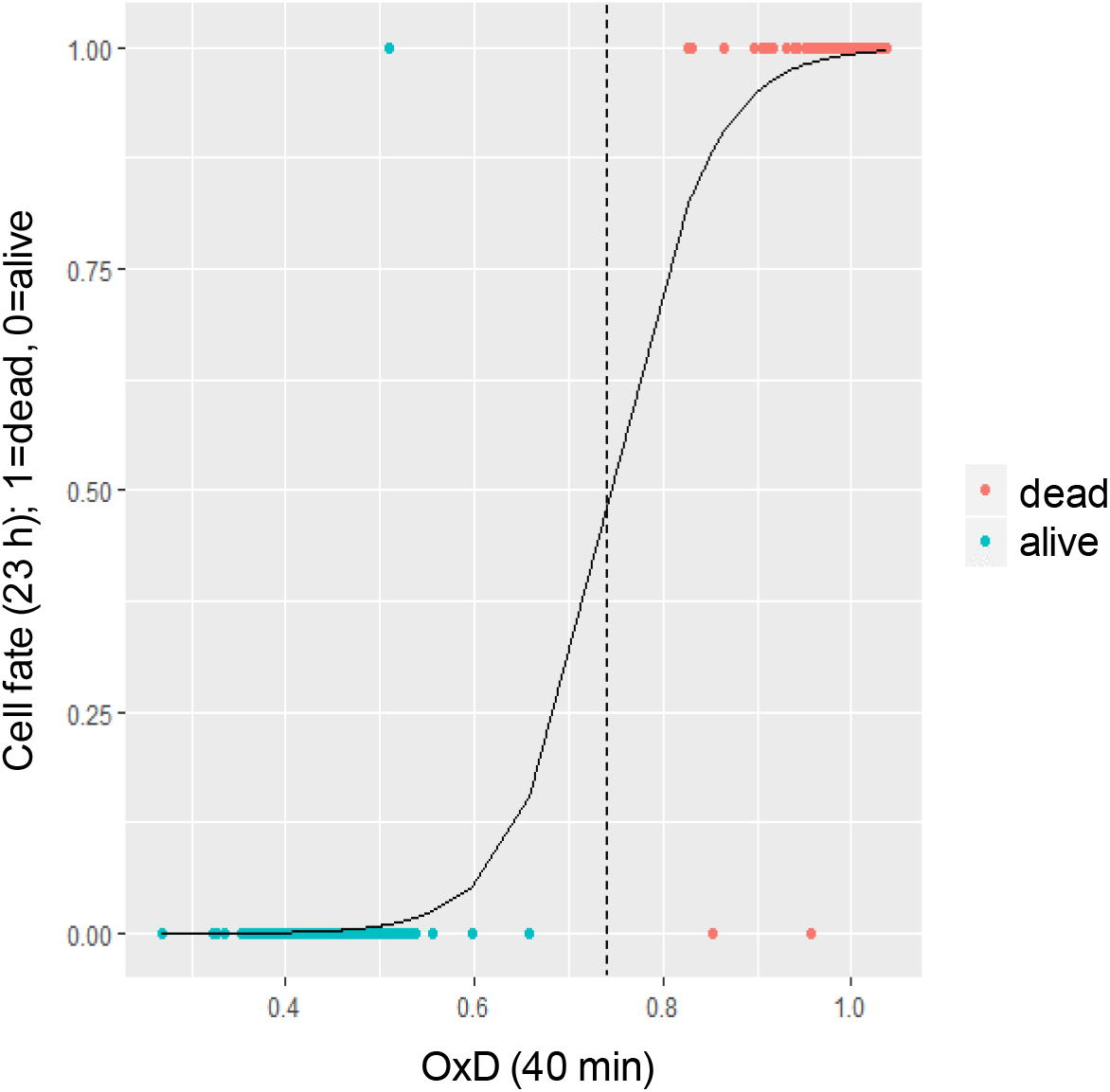
Logistic regression analysis of cell fate vs. chl-roGFP oxidation. Cell fate (1=dead, red; 0=alive, cyan) as measured ∼23 h post 80 μM H_2_O_2_ treatment using Sytox staining vs. chl-roGFP OxD 40 min post treatment. Each dot is data from a single cell, n=252 cells. Logistic regression is shown as a solid line. Threshold of OxD=74% used for cell fate predictions is shown as a dashed line.

**Fig. S9.**
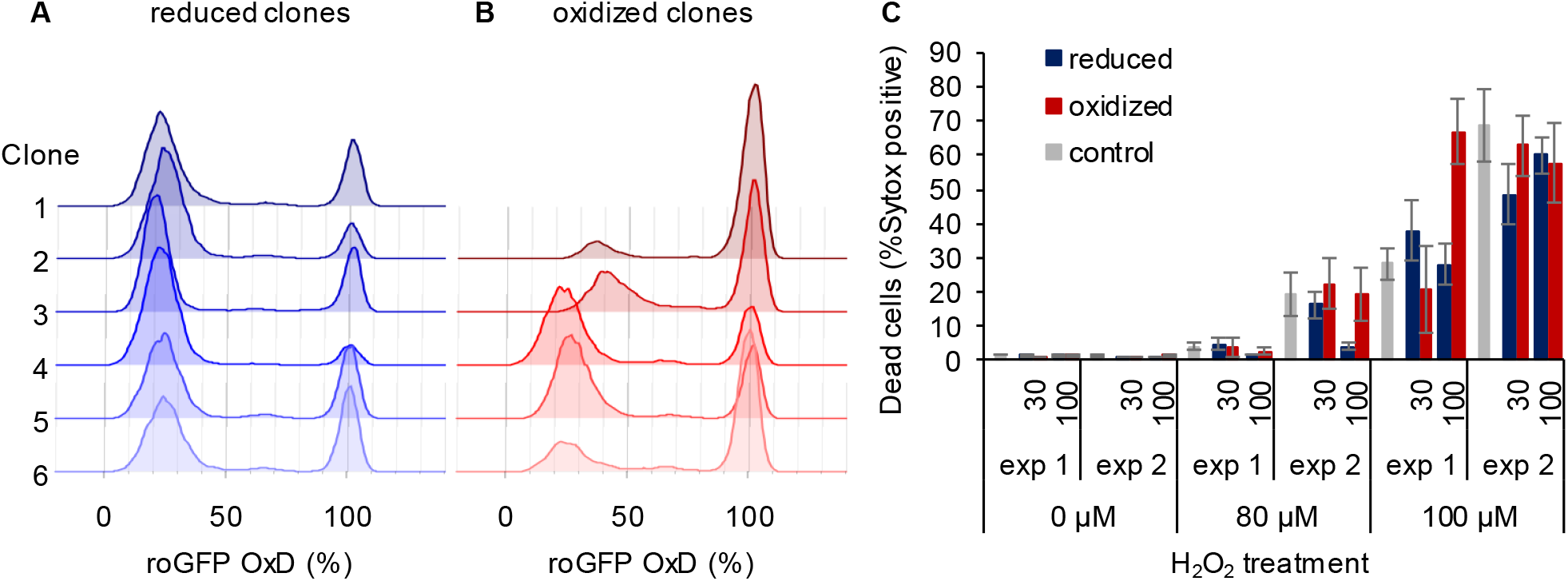
Sorted clonal populations maintain the bi-stable phenotype in chl-roGFP response to H_2_O_2_. (***A-B***) The distribution of chl-roGFP OxD 40-45 min post re-exposure to 80 μM H_2_O_2_ treatment in clonal populations derived from sorted single cells of different origins, 3 weeks post sorting. The “reduced” (A) and “oxidized” (B) subpopulations were sorted 100 minutes post 80 μM H_2_O_2_ treatment based on chl-roGFP oxidation (Fig. 3C). Each histogram is of a single clone, ≥9900 cells per histogram, 6 representative clones per group are shown. The same phenomenon was observed in 2 independent experiments in all examined clones (≥18 clones per group), for visualization data from one is shown. (***C***) The fraction of dead cells 24 h post H_2_O_2_ treatment of clones originating from single cells sorted 30 and 100 minutes post H_2_O_2_ treatment or untreated control, as measured by positive Sytox staining. Data is shown as mean ± SEM, n≥5 per column. Data of 2 independent experiments.

**Fig. S10.**
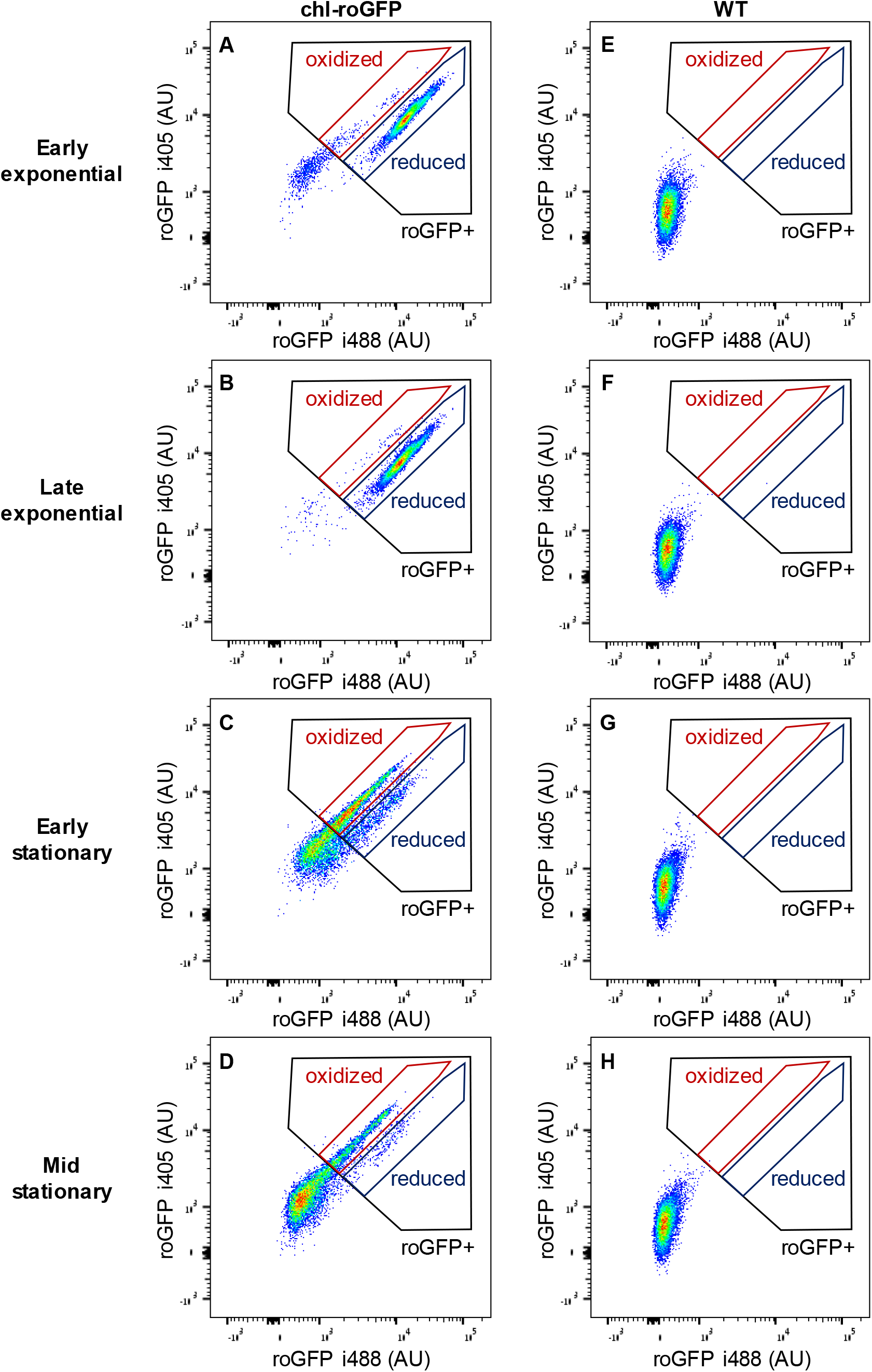
Distinct chl-roGFP subpopulations observed in exponential and stationary cultures. Density plots of fluorescence intensity (AU) of roGFP i405 vs. roGFP i488 in pseudo-color of chl-roGFP (A-D) and WT (no roGFP expression, E-H) strains in early exponential (A, E), late exponential (B, F), early stationary (C, G) and mid stationary (D, H) cultures, 1 h post 50 μM H_2_O_2_. The ratio between i405 and i488 increases upon roGFP oxidation (see methods). Cells with positive roGFP fluorescence (roGFP+, black) and the “oxidized” (red) and “reduced” (blue) subpopulations are marked. The experiment was done in biological triplicates, one representative repeat is shown for visualization. ≥9,000 cells per sample.

**Fig. S11.**
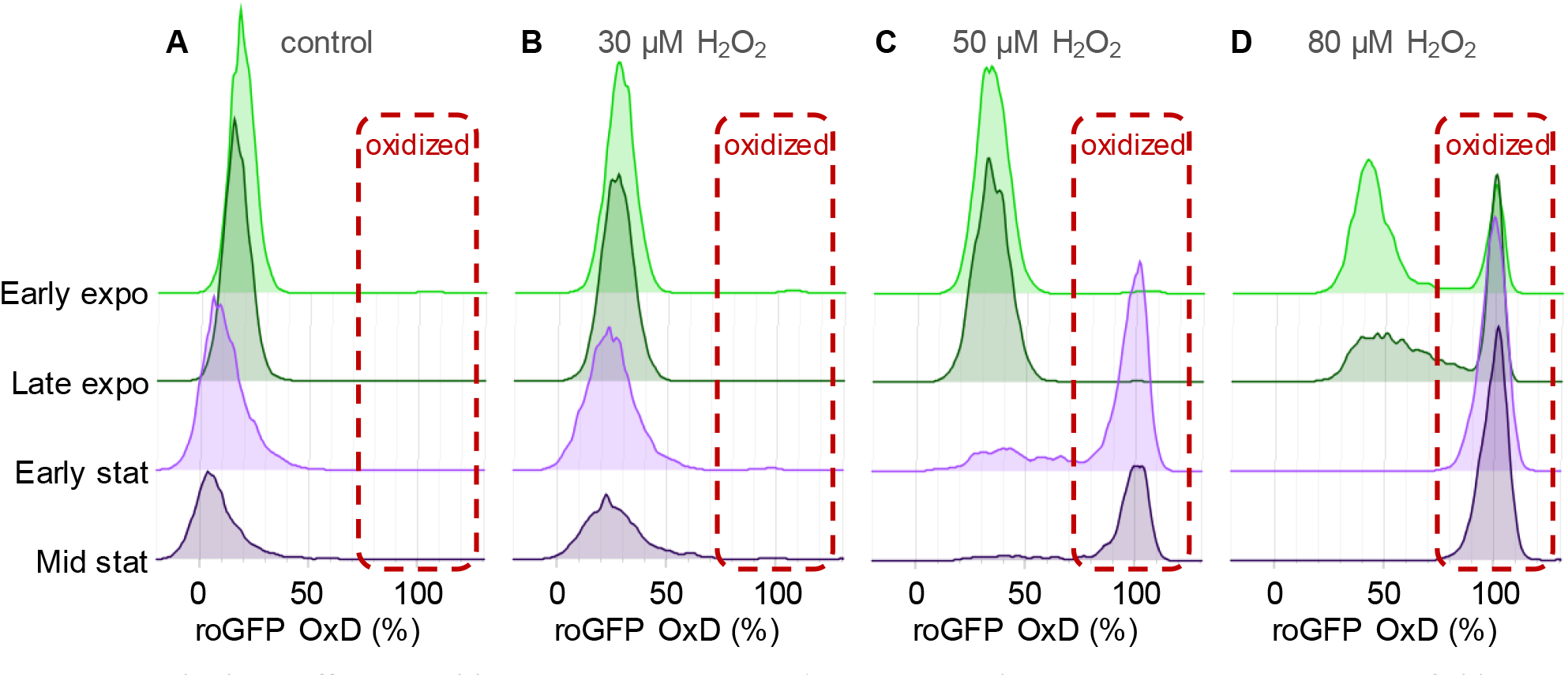
Growth phase effects on chl-roGFP response to oxidative stress. Flow cytometry measurements of chl-roGFP OxD distribution 1 h post treatment with 0 μM (A), 30 μM (B), 50 μM (C) and 80 μM (D) H_2_O_2_ in early exponential (light green), late exponential (dark green), early stationary (light purple), and mid stationary cultures (dark purple). Each histogram consists of >5,000 roGFP+ cells, except mid stationary which consists of >2,000 roGFP+ cells. The experiment was done in biological triplicates, one representative repeat is shown for visualization.

**Fig. S12.**
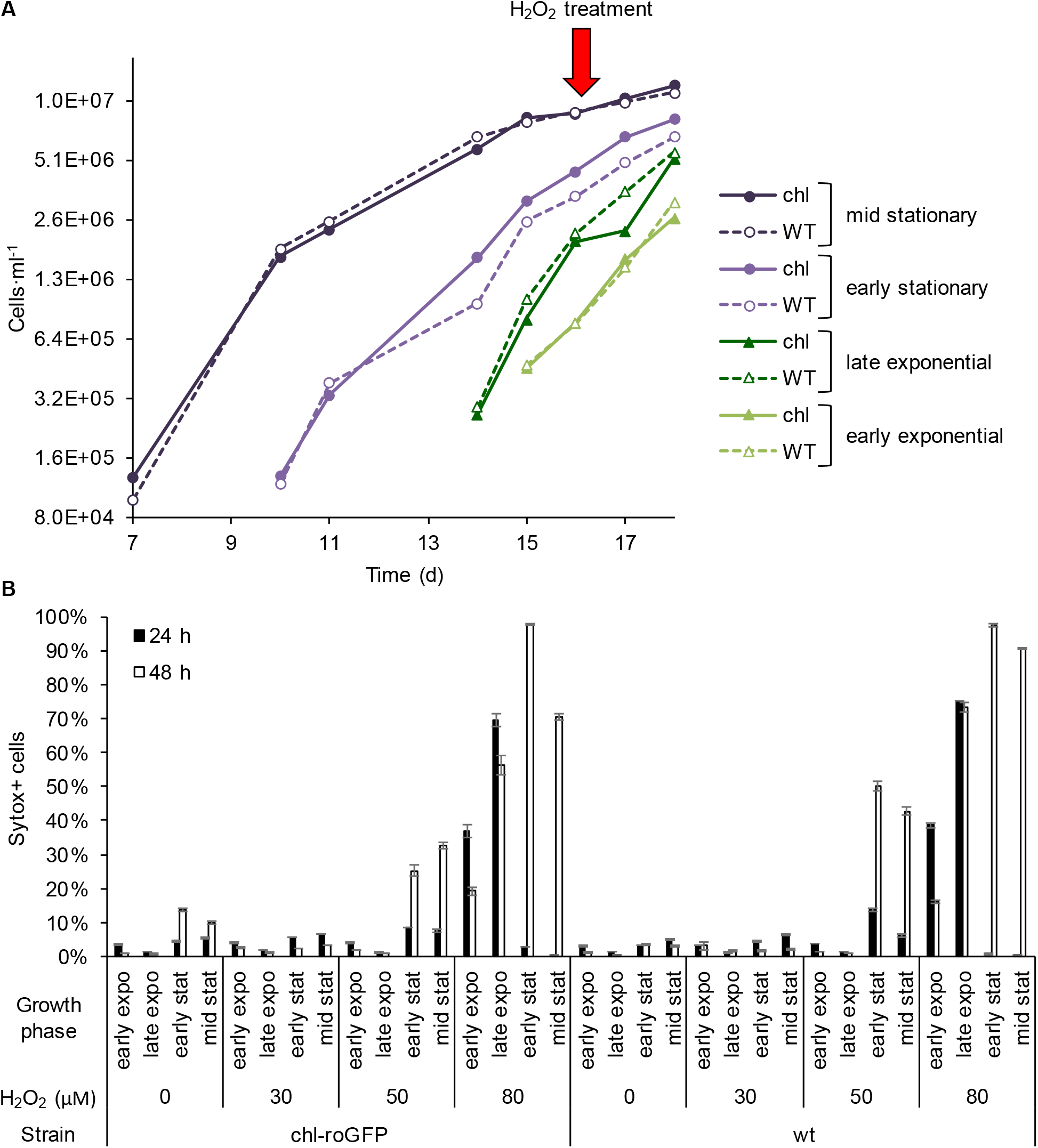
Culture age affects H_2_O_2_ sensitivity and time of death. (***A***) Culture growth curve as measured by flow cytometry in wild type (WT, dashed line) and chl-roGFP strains (chl, same as in Fig. 6 A, solid line). The culture batch was sequentially diluted over time starting at day 0 to generate cultures at different growth phases. Red arrow marks the day of H_2_O_2_ treatment, cell concentration was adjusted prior to treatment. (***B***) The fraction of dead cells 24 h (black) and 48 h (white) post 0-80 μM H_2_O_2_ treatments in cultures at different growth phases as measured by positive Sytox staining in chl-roGFP and wt strains. Mean ± SEM, n=3 biological repeats.

**Fig. S13.**
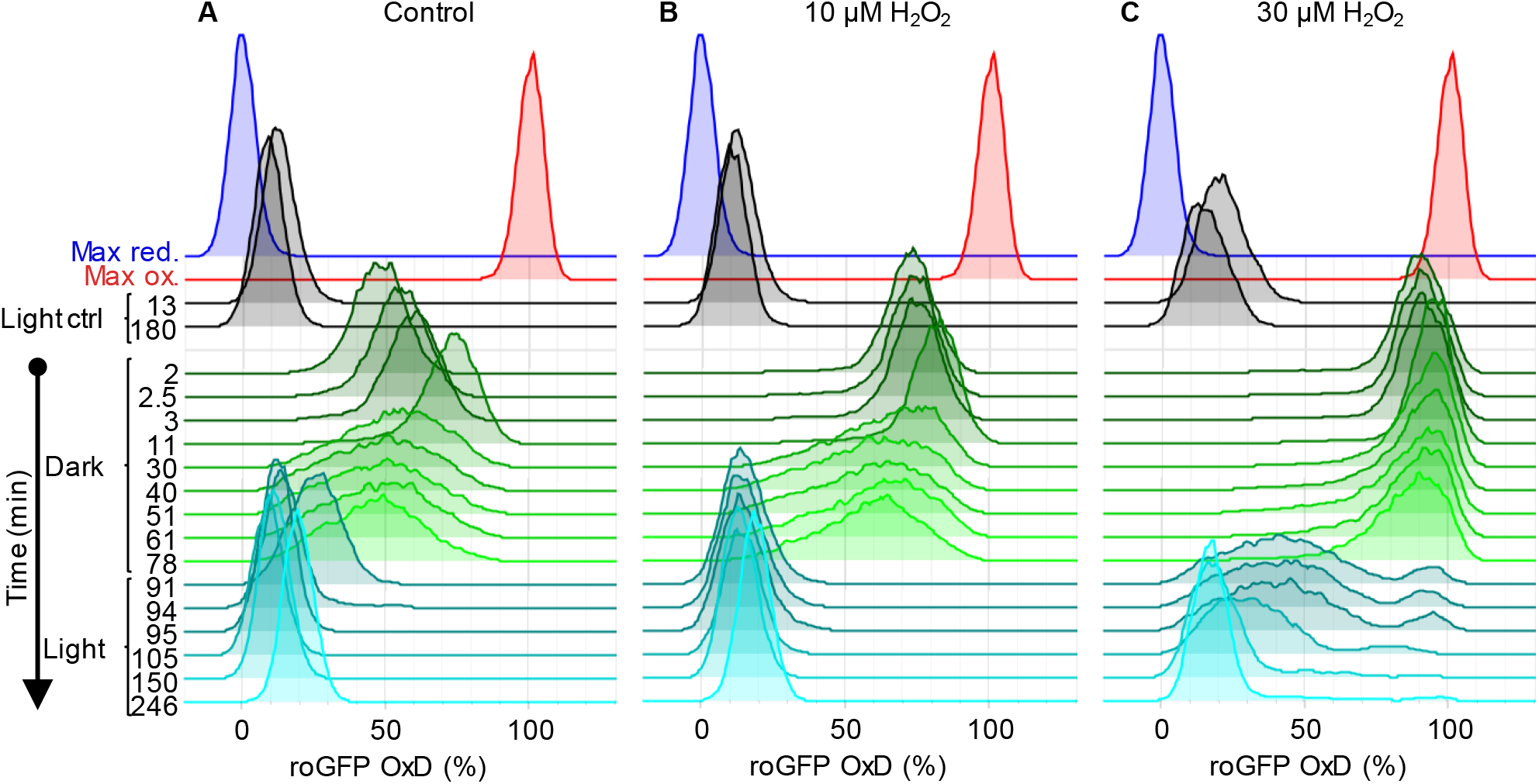
Redox response of chl-roGFP to transition to the dark. Flow cytometry measurements of chl-roGFP oxidation in the population over time. Cells were treated with 0 μM (basal, A, same as in Fig. 2 A), 10 μM (B), and 30 μM H_2_O_2_ (C), and were then transitioned to the dark at time 0 (within 10 minutes post treatment). Cells were kept in the dark for 90 minutes (green) and were then transferred back to the light (cyan). For reference, chl-roGFP OxD following the same H_2_O_2_ treatment but without transition to the dark (black) and following maximum oxidation (200 μM H_2_O_2_, red) and maximum reduction (2 mM DTT, blue) are shown. The experiment was done in triplicates that were highly similar, for visualization purposes the first replica is shown. Each histogram consists of >8000 cells.

**Fig. S14.**
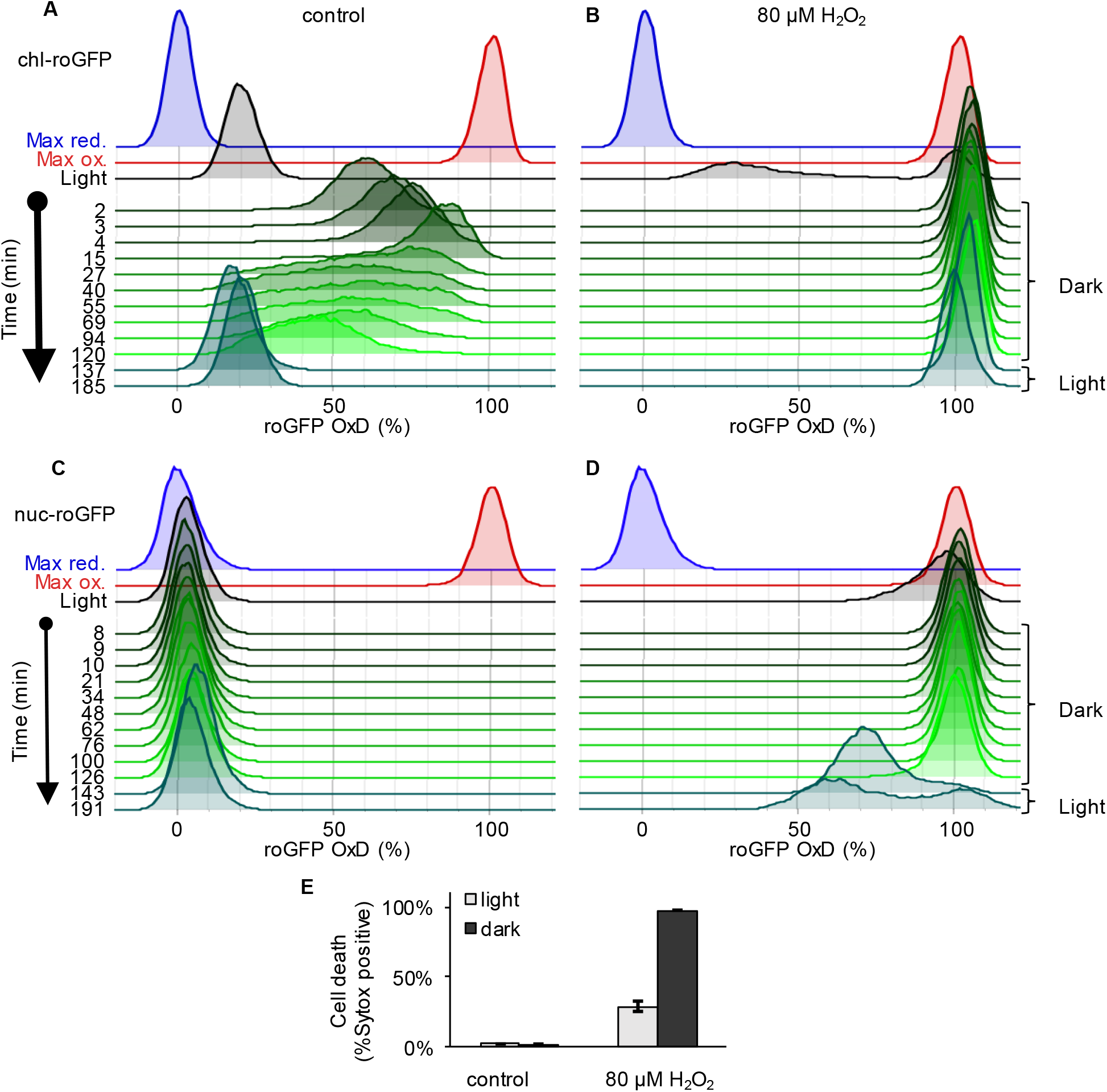
Responses of chloroplast and nucleus targeted roGFP to transition to the dark. Measurements of roGFP oxidation response in the chloroplast and nucleus to a temporal transition to the dark. (***A-D***) Flow cytometry measurements of roGFP oxidation dynamics in the population over time in the chloroplast (A-B) and nucleus (C-D). Cells were treated with 80 μM H_2_O_2_ (C, D) or untreated control (A, C), and were then transitioned to the dark at time 0 (within 4 minutes post treatment). Cells were kept in the dark for 120 minutes (green) and were then transferred back to the light (dark cyan). For reference, roGFP OxD following the same H_2_O_2_ treatment but without transition to the dark (black, ∼130 min post treatment) and following maximum oxidation (200 μM H_2_O_2_, red) and maximum reduction (2 mM DTT, blue) are shown. (***E***) The fraction of dead cells 24 h post H_2_O_2_ treatment with or without transition to the dark (“dark” and “light” respectively) as measured by positive Sytox staining in chl-roGFP strain. Data is shown as mean ± SEM, n=3 biological repeats.

**Fig. S15.**
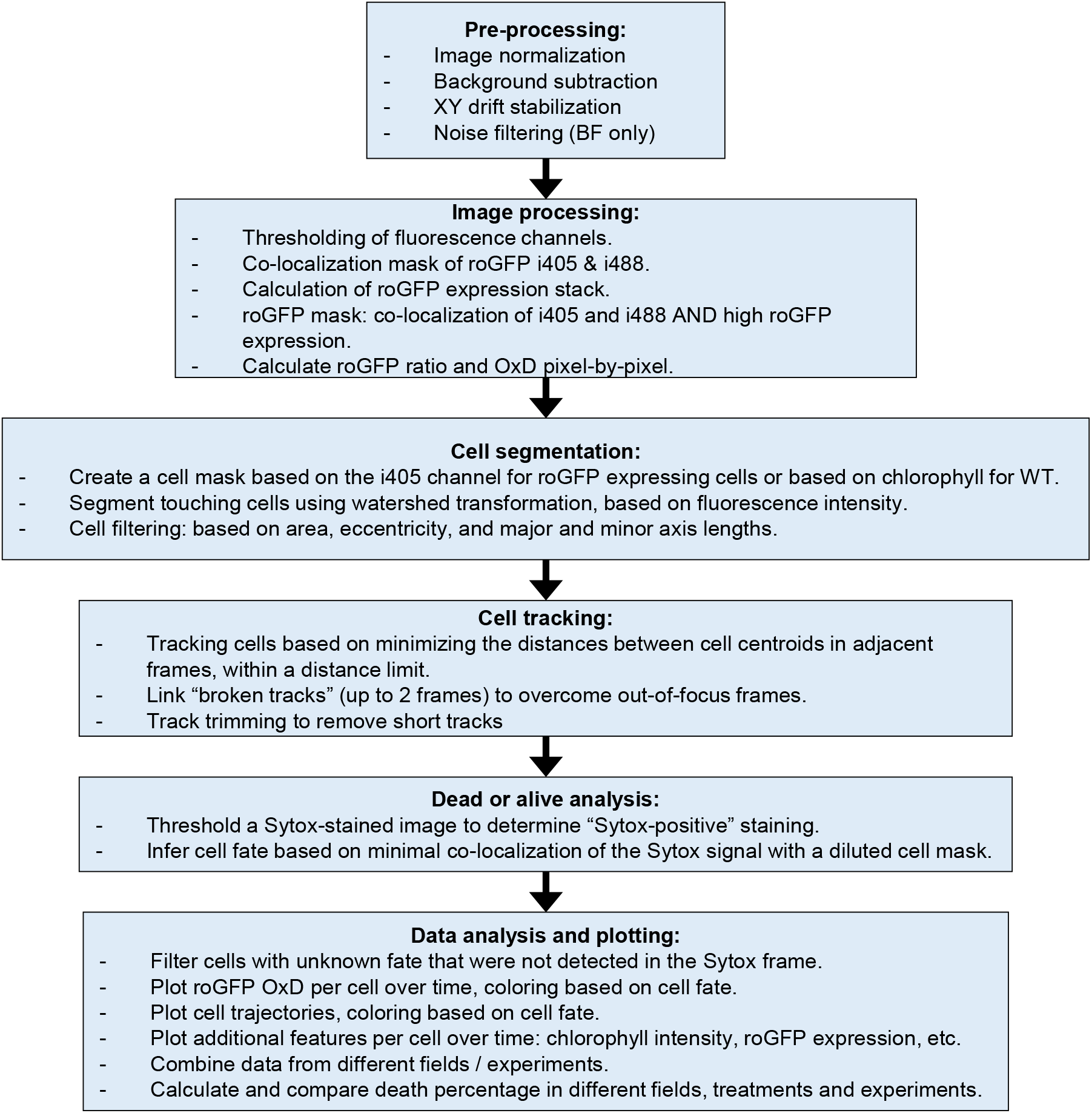
Schematic representation of the microfluidics image analysis pipeline.

**Fig. S16.**
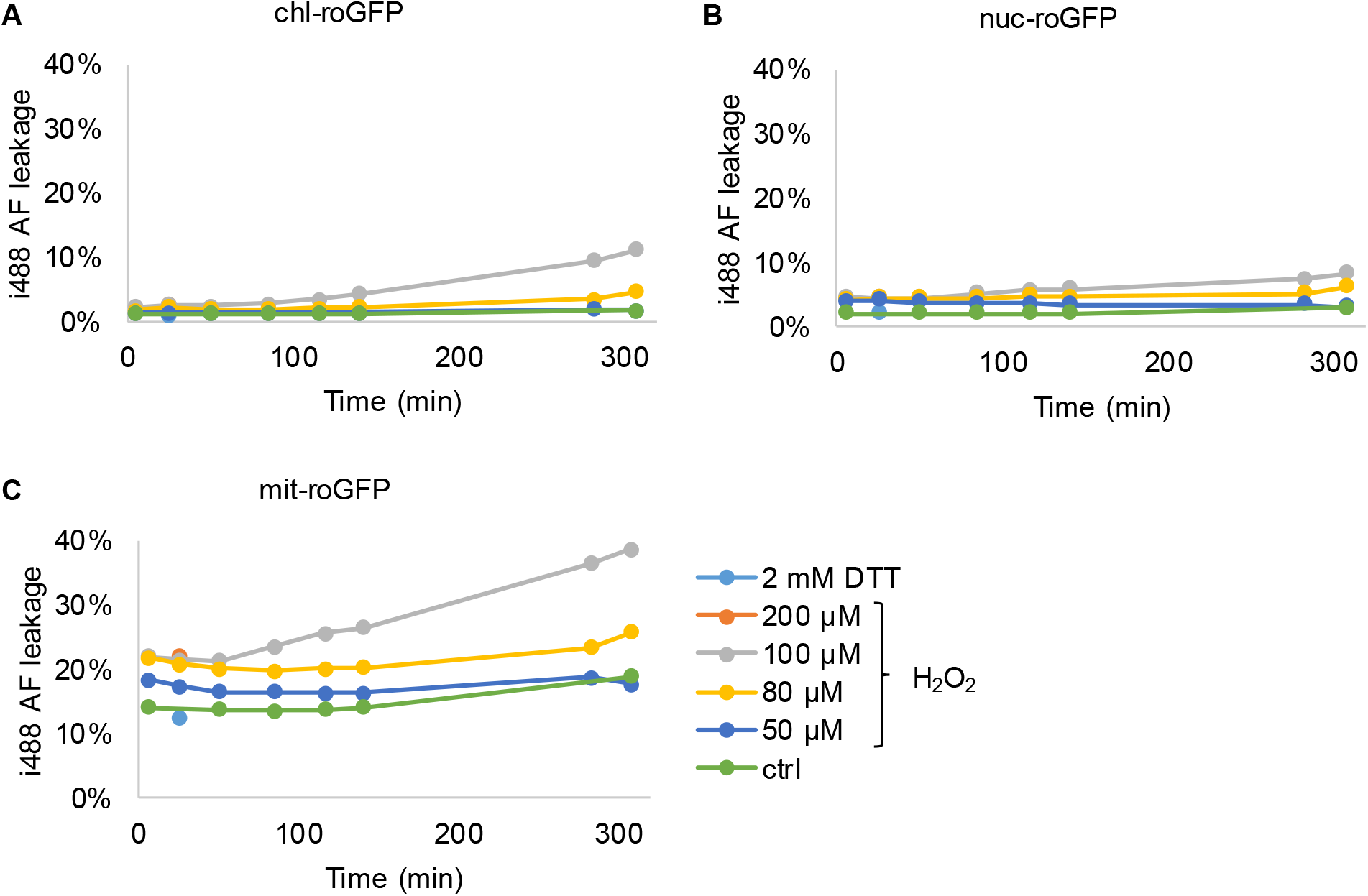
AF leakage into i405 channel in roGFP strains following H_2_O_2_ treatment. Flow cytometry measurements of i405 AF leakage in *P. tricornutum* strains expressing roGFP targeted to the chloroplast (chl-roGFP, A), nucleus (nuc-roGFP, B) and mitochondria (mit-roGFP, C) over time following treatments of 0-200 μM H_2_O_2_ or 2 mM DTT. Leakage into i405 channel was calculated by the mean i405 of the WT strain (n=3 biological repeats) divided by the mean i405 of the roGFP expressing strain (n=3 biological repeats) at the same time-point following the same treatment.

**Fig. S17.**
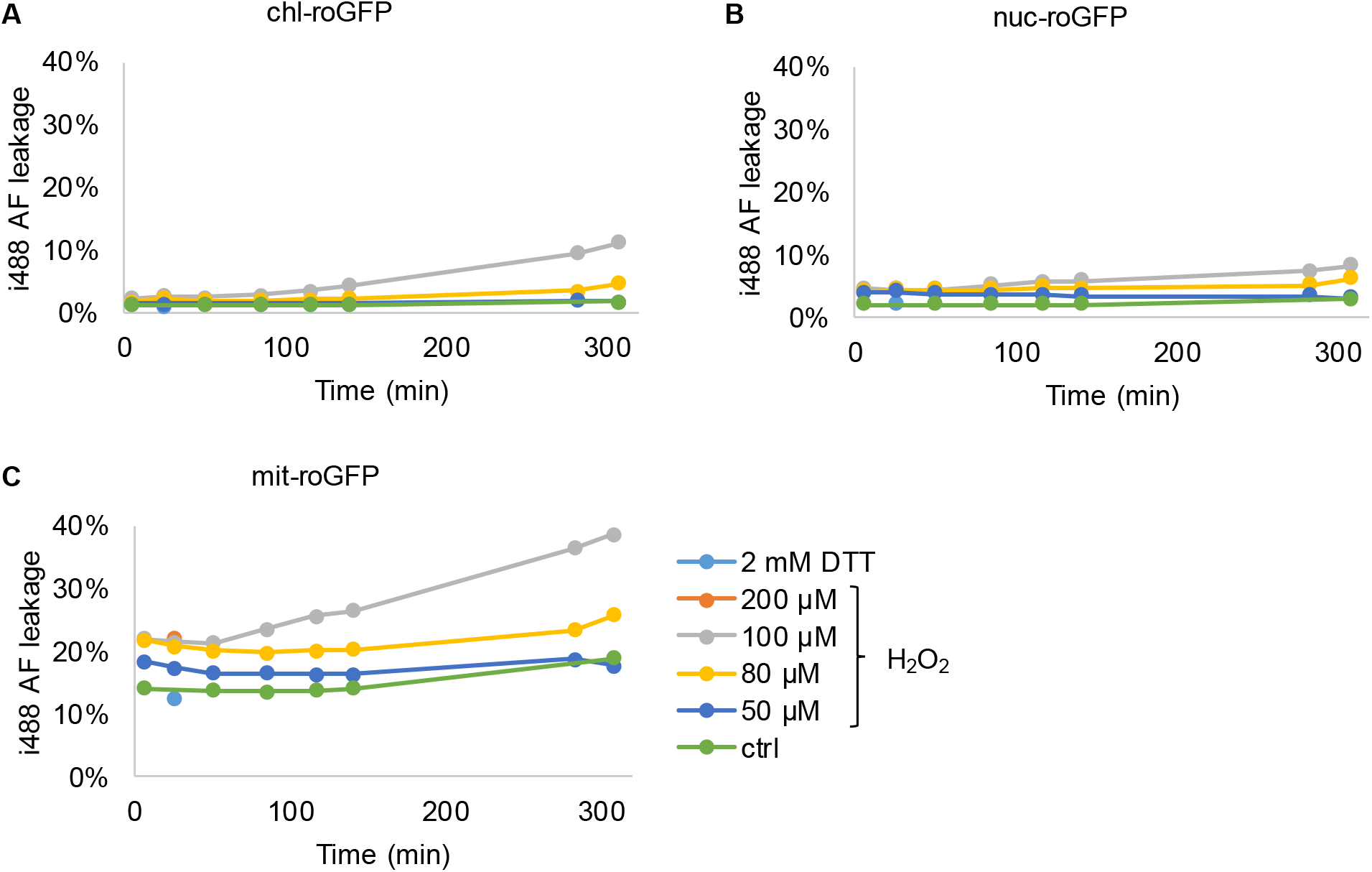
AF leakage into i488 channel in roGFP strains following H_2_O_2_ treatment. Flow cytometry measurements of i488 AF leakage in *P. tricornutum* strains expressing roGFP targeted to the chloroplast (chl-roGFP, A), nucleus (nuc-roGFP, B) and mitochondria (mit-roGFP, C) over time following treatments of 0-200 μM H_2_O_2_ or 2 mM DTT. Leakage into i488 channel was calculated by the mean i488 of the WT strain (n=3 biological repeats) divided by the mean i488 of the roGFP expressing strain (n=3 biological repeats) at the same time-point following the same treatment.

**Table S1.** Growth phase effects on roGFP fluorescence intensity. Flow cytometry measurements of chl-roGFP and WT (no roGFP) strains at different growth phases (see Fig. 6 A). Cell concentration (Cells·ml^−1^) is as measured before cell concentration adjustments. Dynamic range (R_ox_/R_red_) was calculated by dividing the mean roGFP ratio under maximum oxidation conditions (200 μM H_2_O_2_) by that under maximum reduction conditions (2 mM DTT). All other measurements were performed on untreated cultures. Auto-fluorescence (AF) leakage into i488 and i405 channels were calculated by the mean intensity of WT strain divided by the mean intensity of chl-roGFP strain in the i488 and i405 channels respectively. Cells with high AF in the 405ex/450em channel (“high AF cells”) were excluded from analysis (see methods). Population frequencies are shown as mean ± SEM. 10,000 cells analyzed in total per sample, n=3 biological repeats.

| Strain | Growth phase | Cells·ml<sup>-1</sup> | roGFP+ freq. | roGFP- freq. (low AF) | High AF cells freq. | Mean roGFP OxD (steady state) | Mean i488 AF leakage | Mean i405 AF leakage | roGFP dynamic range | roGFP+ cells per sample (min-max) |
| --- | --- | --- | --- | --- | --- | --- | --- | --- | --- | --- |
| chl-roGFP | early expo. | 7.8·10<sup>5</sup> | 93.3±0% | 6.6±0% | 0.1±0% | 19.3±0.2% | 1.7% | 6.5% | 4.96 | 9388-9396 |
|  | late expo. | 2·10<sup>6</sup> | 99.1±0% | 0.8±0% | 0.1±0% | 16.5±0.2% | 2.1% | 7.6% | 4.9 | 9935-9949 |
|  | early stat. | 4.5·10<sup>6</sup> | 53.6±0.2% | 43.3±0.3% | 3.1±0.2% | 12±0.2% | 3.8% | 13.6% | 4.57 | 5380-5449 |
|  | mid stat. | 8.8·10<sup>6</sup> | 25.1±0.3% | 72.6±0.3% | 2.3±0.1% | 9.5±0.2% | 3.4% | 12.8% | 4.53 | 2462-2575 |
| WT | early expo. | 7.7·10<sup>5</sup> | 0±0% | 100±0% | 0±0% | - | - | - | - | - |
|  | late expo. | 2.2·10<sup>6</sup> | 0±0% | 99.9±0% | 0±0% | - | - | - | - | - |
|  | early stat. | 3.4·10<sup>6</sup> | 0±0% | 99±0% | 1±0.1% | - | - | - | - | - |
|  | mid stat. | 9·10<sup>6</sup> | 0±0% | 98±0% | 2±0% | - | - | - | - | - |

**Table S2.** Dynamic range of roGFP targeted to different organelles. The dynamic range (R_ox_/R_red_) was calculated by dividing the mean roGFP ratio under maximum oxidation conditions (200 μM H_2_O_2_) by that under maximum reduction conditions (2 mM DTT) using data from flow cytometry and microfluidics imaging experiments.

| *P. tricornutum* strain | Flow cytometry<br>dynamic range | Microfluidics imaging<br>dynamic range |
| --- | --- | --- |
| chl-roGFP | 5.57 | 5.42 |
| nuc-roGFP | 5.94 | - |
| mit-roGFP | 3.50 | - |

### Movie legends

**Movie S1. Microfluidics *in-vivo* imaging of chl-roGFP oxidation over time following H_2_O_2_ treatment**. Oxidation of chl-roGFP cells was imaged over time using a customized microfluidic setup and epifluorescence microscopy with controlled flow, light and temperature conditions (SI appendix). Movie of chl-roGFP OxD in pseudo-color of cells treated with 80 μM H_2_O_2_ at time 0, the first frame is before treatment. Time stamp represents time post treatment (hh:mm). Color bar as in Fig. 5.

**Movie S2. Microfluidics *in-vivo* imaging of basal chl-roGFP oxidation over time**. Oxidation of chl-roGFP cells was imaged over time using a customized microfluidic setup and epifluorescence microscopy with controlled flow, light and temperature conditions (SI appendix). Movie of chl-roGFP OxD in pseudo-color of cells treated with fresh media (control) at time 0, the first frame is before treatment. Time stamp represents time post treatment (hh:mm). Color bar as in Fig. 5.

